# Unified high-resolution immune cell fraction estimation in blood tissue from birth to old age

**DOI:** 10.1101/2025.02.02.636167

**Authors:** Xiaolong Guo, Mahnoor Sulaiman, Alexander Neumann, Shijie C Zheng, Charlotte AM Cecil, Andrew E Teschendorff, Bastiaan T Heijmans

## Abstract

**Background:** Blood is composed of many different immune cell-types, whose proportions vary throughout life. If not accounted for, these variations can seriously cause confounding or hamper interpretation of DNAm-based biomarkers. Although cell-type deconvolution methods can address this challenge for cord and adult blood, there is currently no method that can be applied to blood tissue from other age groups, including infants and children.

**Results:** Here we construct and extensively validate a DNAm reference panel, called UniLIFE, for 19 immune cell-types, applicable to blood tissue of any age. We use UniLIFE to delineate the dynamics of immune-cell fractions from birth to old age, and to infer disease associated immune cell fraction variations in newborns, infants, children and adults. In a prospective longitudinal study of type-1 diabetes in infants and children, UniLIFE identifies differentially methylated positions that precede type-1 diabetes diagnosis and that map to diabetes related signaling pathways. In contrast to previous studies, we are able to validate these biomarker associations in an independent DNAm dataset comprising purified monocytes from monozygotic twins discordant for type-1 diabetes, but not in lymphocytes, highlighting the importance of epigenetic changes in the innate immune system in the development of type-1 diabetes.

**Conclusions:** In summary, we present a life course immune-cell estimator for blood tissue of any age to help improve the identification and interpretation of blood-based DNAm biomarkers for any age groups and specially for longitudinal studies that include infants and children. The UniLIFE DNAm reference panel and algorithms to estimate cell-type fractions and perform cell-type deconvolution are available from our EpiDISH Bioconductor R-package: https://bioconductor.org/packages/release/bioc/html/EpiDISH.html

## Background

DNA methylation (DNAm) is an epigenetic mark that plays an important role in gene regulation, aging and disease ^1^. Like gene expression, DNAm differs significantly between cell-types ^2–5^. Due to the high cost and time consuming task of cell-sorting, especially when implemented at large scale (e.g. as in the case of population-based studies), epigenome studies typically examine DNAm in bulk tissue, which comprises a mixture of many different cell-types ^1^. Thus, DNAm measurements in bulk-tissue are subject to potential confounding by variations in cell-type composition ^6–8^. To address this challenge, cell-type deconvolution algorithms and DNAm reference panels have been developed ^9–13^ that allow researchers to estimate cell-type fractions, which are necessary to adjust for the underlying cell-type heterogeneity, thus circumventing the need to experimentally measure the actual cell-type composition.

One of the most commonly used bulk tissues for epigenetic studies is blood, including cord blood at birth ^14–21^ and whole/peripheral blood at later ages ^22^. While both involve ‘blood’, there are key differences between these tissue types, with cord blood containing a unique set of cell types that rapidly decline after birth, e.g. nucleated red blood cells (nRBC), as well as cell-types that have different properties compared to their adult counterparts, for example the degree of ‘naivety’ of B or T cells ^23^. To capture these differences, distinct cell-type estimation panels, called DNAm reference matrices/panels, have been created ^10,24–28^. The Gervin DNAm reference matrix, which estimates fractions for seven major cell groups in cord-blood (B cells, Granulocytes, Natural Killer (NK) cells, Monocytes, CD4T, CD8T, and nRBCs), is considered the gold standard for studies using DNAm data at birth ^25^. For adult blood, the Salas ^27^ and updated EpiDISH ^29^ DNAm reference matrices estimate fractions for 12 immune cell types. While both types of panels are effective within their specific contexts, i.e. cord-blood/birth and adulthood, respectively, the use of separate DNAm reference panels for different life stages can pose key challenges for the field. In particular, any analysis aiming to investigate longitudinal dynamics or epigenetic timing effects using blood at different time points, say cord-blood and mid-age, would need to apply different cell-type estimation panels, which could cause confounding between tissue composition and epigenetic changes that happen in individual cell-types ^30^. Moreover, current cord and adult-blood reference panels are not directly applicable to blood sampled in between these ages, say from infants, children or young adolescents, since in the early life stages, immune cells are undergoing maturation with the corresponding fractions also displaying highly variable dynamics ^23,31–33^. As such, the development of a ‘universal’ or ‘lifecourse’ blood DNAm reference matrix would enable researchers to use the same panel at any time point, thus reducing confounding between timepoints, and enhancing the reliability and interpretability of DNAm-based biomarkes ^34^ ^35^. Given the increasing number of cohort studies with DNAm measured repeatedly across time, specially in infants, children and adolescents ^31,36–39^, there is an urgent need to develop such a lifecourse DNAm reference panel.

The purpose of this study is to create and extensively validate a novel, unified ‘lifecourse’ DNAm reference panel, called UniLIFE, that can be applied to blood samples of any age, from birth onwards, including infancy, childhood, adolescence and adulthood. Our validation is performed on (i) independent sorted samples from cord and adult blood derived with two different technologies (Illumina beadarrays and whole-genome bisulfite sequencing (WGBS)), (ii) on unsorted cord and adult blood samples with matched cell-counts, and (iii) on blood samples from infants, children and adolescents. We use our lifecourse blood DNAm reference panel to delineate the dynamic changes in blood composition that occur immediately after birth and during the first months and years of life, a time period for which current panels are not directly applicable and where we also currently lack much knowledge ^40^. Indeed, charting the dynamic landscape of immune-cell composition in blood during the earliest life-stages is critically important as this can also shed light on future disease predisposition or indicate compositional changes that are diagnostic of disease ^40,41^. In line with this, we demonstrate that our reference panel can capture disease-associated cell-type composition variations during the first years of life, including DNAm changes that predate the onset of type-1 diabetes and which map to relevant signaling pathways.

### Implementation

The UniLIFE DNAm reference panel is a matrix of DNAm values defined over 1906 marker CpGs and 19 immune cell-types. Calling this panel “*centUniLIFE.m*”, and given an independent DNAm dataset matrix *X* defined over CpGs (Illumina 450k, Illumina EPIC1/2 or WGBS) and samples, we estimate fractions for the 19 immune cell-types using the following pseudocode in R-programming language:

➢ *library(EpiDISH)*
➢ *data(centUniLIFE.m)*
➢ *estF19CT.m <-epidish(X, cent=centUniLIFE.m, method=”RPC”, maxit=500)$estF* where *estF19CT.m* is a matrix containing the estimated immune cell fractions for samples (rows) and each of the 19 immune cell-types. With these estimated fractions it is then possible to (i) identify disease associated differentially methylated cytosines (DMCs) that are not driven by changes in cell-type composition, and (ii) to identify cell-type specific DMCs, as described by us previously ^42^.

## Results

### An axis of DNAm variation associated with cell maturity and age

Our overall strategy to build a universal DNAm reference matrix for blood of any age is depicted in **Fig.1a.** Briefly, we collected Illumina DNAm data from 7 sorted cord-blood ^25,43–45^ and 12 sorted adult immune cell types ^27^, and subjected these datasets to a stringent quality control and normalization procedure, removing samples of low purity (**Methods, SI figs.S1-S2**) ^29^. This resulted in an integrated DNAm dataset comprising 375,506 CpGs and 220 sorted samples, encompassing 19 immune cell subtypes, 7 from cord-blood and 12 from adult blood (**Fig.1a**). PCA-analysis on this merged dataset revealed broad segregation of samples by cell-type, with PC-1 describing a myeloid vs lymphoid differentiation axis, and with PC-2 describing an axis of variation reflecting cell-maturity (**Fig.1b**). Supporting the interpretation of PC-2 as a maturity axis, we note that along this axis, adult naïve B and T-cells are more similar to their cord-blood counterparts than to adult memory B and T-cells.

**Figure-1:**
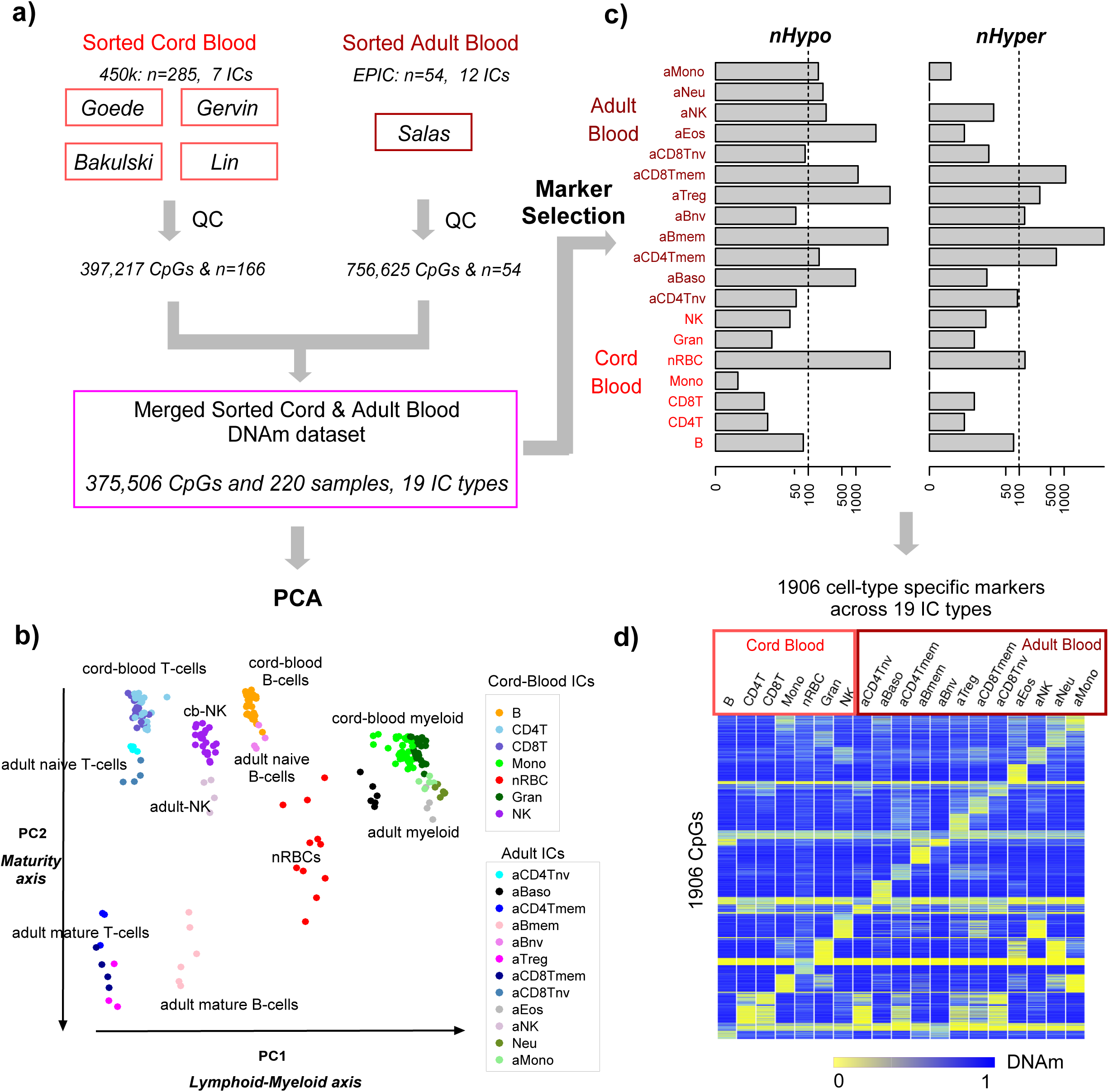
Construction of the unified cord adult blood DNAm reference matrix. **a)** Four independent cord blood Illumina 450k datasets encompassing 7 immune cell types were processed, underwent separate quality control (QC) analyses which removed samples of low purity. The resulting high quality samples were merged together with an Illumina EPIC DNAm dataset of 12 sorted adult immune cell types, resulting in an integrated DNAm dataset of 220 sorted samples encompassing 19 immune cell-types. **b)** PCA scatterplot (PC1 vs PC2) of this merged dataset with samples annotated by one of the 19 immune cell-types. Specific clusters of cells are labeled in the plot to guide the reader. **c)** Barplot contrasting the number of cell-type specific markers for each of the 19 immune cell-types, stratified by whether the marker CpG is hypomethylated or hypermethylated in the given cell-type relative to all other cell-types. The x-axis labeling the numbers of markers are displayed on a log-scale. Based on this distribution, a total of 1906 cell-type specific markers were selected. **d)** Heatmap representation of the resulting unified DNAm reference matrix for blood encompassing 1906 cell-type specific marker CpGs and 19 immune cell types.

Moreover, for cells from the innate immune system (myeloid and NK-cells), the difference between cord and adult cells was much smaller than that between cord-blood and mature lymphocytes. The PCA plot further revealed that all 19 immune cell-types are separate entities, and that adult naïve T and B-cells are distinguishable from their cord-blood counterparts. Overall, this data indicates that whilst PC-2 is predominantly reflecting a maturity axis, this same axis of variation also discriminates aged naïve/immature lymphocytes from naïve/immatures lymphocytes found in cord-blood, suggesting that PC-2 is simultaneously also an axis of ‘aging’.

### Construction of a 19 immune cell type DNAm reference matrix from cord and adult blood

Next, we devised a statistical method to select cell-type specific markers, supplementing an empirical Bayes method for detecting differentially methylated cytosines (DMCs) with a novel gap-specificity score and triangulation principle, aimed at maximizing cell-type specificity of the markers (**Methods, SI fig.S3-4**). Given that the most specific regions for adult cell-types are unmethylated in the given adult cell-type ^3^, we first aimed to identify at least 100 cell-type specific hypomethylated markers per cell-type. For most adult mature cell-types, we could identify at least 100 cell-type specific hypomethylated markers, consistent with Loyfer et al ^3^, but for adult naïve and cord-blood cell subtypes, this was not the case, indicating the need to also consider cell-type specific hypermethylated markers (**Fig.1c**). To ensure approximately 100 markers for every cell-type, we also relaxed the gap-specificity score criterion, selecting a number of markers purely based on average fold-changes (**Methods**). This resulted in a total of 1906 marker CpGs across all 19 immune cell subtypes. Finally, a DNAm reference matrix was built by taking the median DNAm value across samples of the same cell-type (**Fig.1d**). We refer to this DNAm reference panel as the UniLIFE (Unified Lifecourse Immune Fraction Estimator) DNAm reference matrix.

### Validation in independent sorted and mixed cord blood datasets

To validate the 19 immune cell-type UniLIFE panel, we first assessed its performance in independent cord-blood DNAm datasets. In an independent 450k dataset of 4 sorted cord-blood subtypes, UniLIFE in conjunction with EpiDISH ^10^ (**Methods**) predicted high purity for each of these four subsets (**Fig.2a**). Only nRBCs displayed a non-negligible fraction of cord-blood granulocytes, potentially indicating a contamination effect (**Fig.2a**). We next applied UniLIFE to sorted cord-blood immune cell samples from IHEC, which were generated with whole-genome bisulfite sequencing. UniLIFE correctly predicted the cell-type of these sorted samples, with the predictions indicating relatively high purity (**SI fig.S5**). Importantly, adult blood cell-types were correctly predicted to have zero or near-zero fractions. Upon generating 100 in-silico mixtures, estimated fractions displayed excellent correlation with the true weights (**Fig.2b**). Next, we applied our DNAm reference matrix to an independent EPIC DNAm dataset of experimental mixtures of 6 cord blood cell subtypes, where the true mixing fractions are known (**Methods**), revealing once again an excellent correlation as well as low root mean square errors (RMSE) (**Fig.2c**). Only for monocytes was the correlation weaker, which we can attribute to the lower dynamic range of mixing weights used for this particular cell-type. Finally, we applied our DNAm reference matrix to an independent cord-blood 450k DNAm dataset with matched FACS cell counts, including nRBCs. Correlations were strong to very strong (**Fig.2d**), except for CD8+ T-cells, which we can attribute to the very low dynamic range of CD8+ T-cell fractions (all FACS counts < 0.08). Indeed, we note that the RMSE for CD8+ T-cells was in fact quite low, suggesting that the lack of correlation is mainly due to the low dynamic range. Importantly, for the nRBC fraction we obtained a reasonably good R and RMSE values. Thus, overall, our DNAm reference matrix can reliably predict fractions in cord blood.

**Figure-2:**
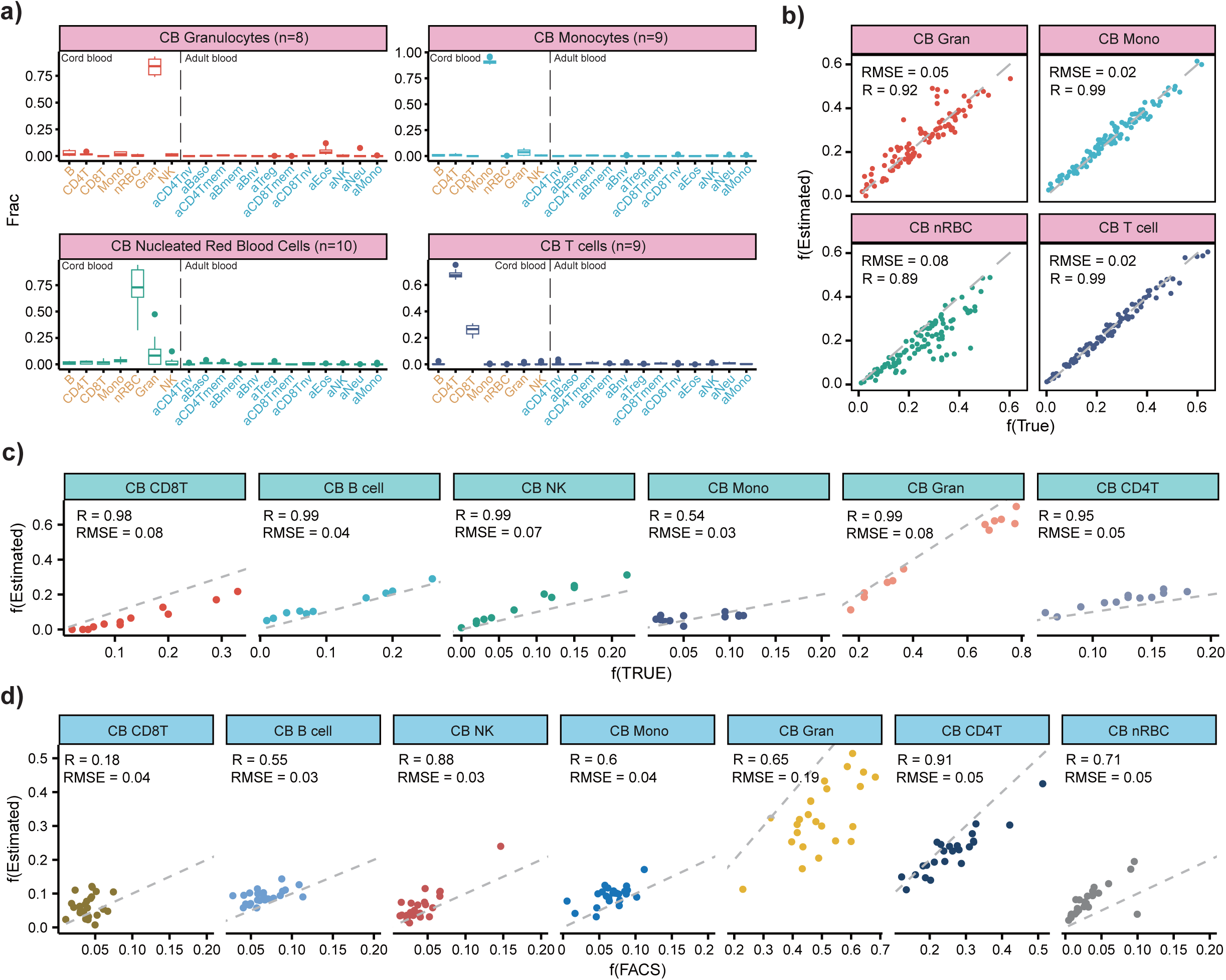
Validation of UniLIFE in cord blood. **a)** Boxplots display the inferred cell type proportions for four sorted cord blood cell types using an independent Illumina 450k dataset of deGoede. The number of sorted samples corresponding to each cell type is provided. **b)** Scatterplots of true mixture weights (x-axis) vs estimated fractions (y-axis) for 100 in-silico mixtures. Each mixture is a linear combination of 4 sorted cord blood samples from a) with weights drawn randomly from a uniform distribution. Pearson R-value and RMSE are given. **c)** Scatterplots of true fractions vs estimated fractions for 6 cord blood cell subtypes using an independent EPIC DNAm dataset of 12 umbilical cord blood experimental mixtures where the underlying mixing proportions were known. For each estimated cell type, we display the Pearson R-value and RMSE. **d)** Scatterplots of FACS-derived fractions vs estimated fractions for 24 independent cord blood 450k DNAm profiles with matched FACS cell-counts for 7 cell-types, as shown. Pearson R-value and RMSE are given.

### Validation in independent sorted and mixed adult blood datasets

Next, we applied a similar strategy to DNAm profiles of adult blood cell-types, experimental mixtures and adult blood with matched experimental cell-counts (**Methods**). We assembled a large collection of independent Illumina 450k and whole genome bisulfite sequencing (WGBS) datasets from adult immune cell subtypes, on which UniLIFE correctly predicted the corresponding cell-types (**Fig.3a**). Of note, the 7 cord-blood cell subtypes obtained zero or near-zero fractions across all these adult samples, as required (**Fig.3a**). To further demonstrate how our DNAm reference matrix can reliably estimate fractions in WGBS data, we generated 100 in-silico mixtures of 10 different WGBS adult blood cell subtypes, in each case, revealing an excellent correlation (**Fig.3b**). To demonstrate validity on EPIC DNAm data, we compared estimated fractions to experimental mixtures of 10 adult blood cell subtype EPIC DNAm profiles, where the true fractions are known, revealing once again very strong correlations and low RMSEs (**Fig.3c**). Finally, in an independent EPIC DNAm dataset of whole blood samples with matched FACS counts, we also observed very strong correlations and low RMSEs despite many fractions displaying a relatively small dynamic range (**Fig.3d**). Overall, these results demonstrate that our 19 immune cell-type DNAm reference matrix can reliably estimate fractions in adult blood.

**Figure-3:**
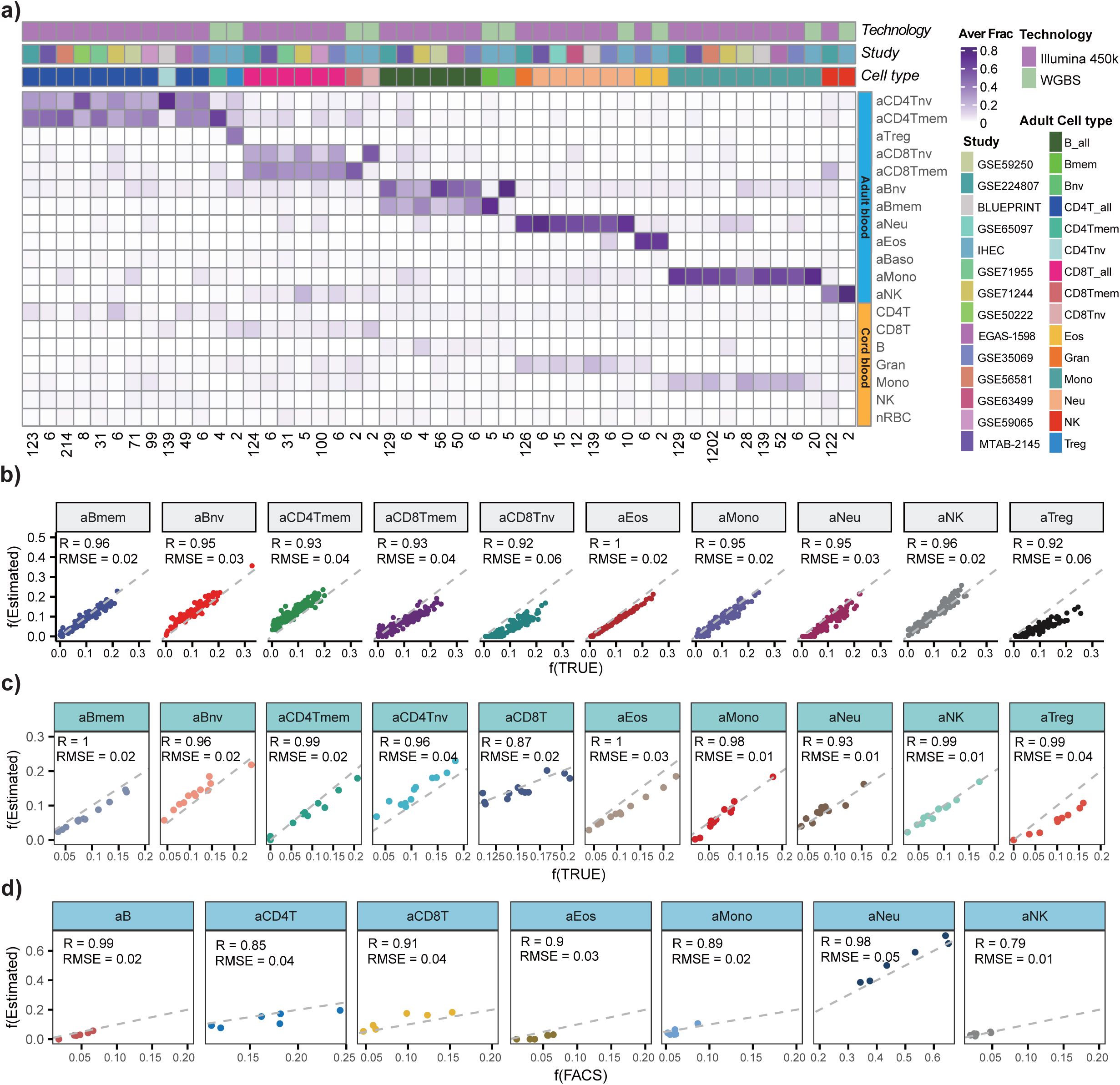
Validation of UniLIFE in adult blood. **a)** Heatmap displays the estimated fractions of sorted samples for each of 12 adult immune-cell subtypes in UniLIFE, as well as the total CD4 + T-cell, total CD8 + T-cell, and total B-cell fractions. The immune-cell type of the sorted sample is indicated by the color bar on top of the heatmap. The technology used to generate the DNAm data of the sorted sample and the study from which the sorted sample derives from are also indicated. The color bar on the right distinguishes between adult (blue) and cord blood (yellow) immune cell types in the UniLIFE reference matrix. The estimated fractions in the heatmap are mean values taken over biological replicates of the cell-sorted samples, with the number of corresponding biological replicate samples indicated at the bottom. **b)** Scatterplots of true mixture weights (x-axis) vs estimated fractions (y-axis) for 100 in-silico mixtures of WGBS DNAm profiles from IHEC. Each mixture is a linear combination of 10 adult WGBS immune cell sorted samples with weights drawn randomly from a uniform distribution. Pearson R-value and RMSE are given. **c)** Scatterplots of true fractions vs estimated fractions for 10 adult blood cell subtypes using the EPIC DNAm data from 12 experimental mixtures where the underlying mixing proportions were known. For each estimated cell type, we display the Pearson R-value and RMSE. **d)** Scatterplots of FACS fractions vs estimated fractions for 6 whole blood EPIC DNAm profiles with matched FACS cell-counts for 7 cell-types, as shown. Pearson R-value and RMSE are given.

### Validation in independent blood datasets of any age

Next, we aimed to demonstrate that UniLIFE can also predict fractions in more complex mixtures composed of cord and adult blood cell-types. This could be relevant for instance in blood of infants or children, where a given leukocyte proportion, say CD4+ T-cells could be modelled as a mix of an immature youthful phenotype, for which those derived from cord-blood could be representative, and an “aged” phenotype encompassing both naïve and mature fractions, as found in adults. We note that this interpretation is well supported by our previous PCA analysis, revealing how PC2 is capturing a maturity axis which is distinct from age, since the adult naïve T-cells cluster more closely to the cord-blood counterparts (**Fig.1b**). Thus, we generated in-silico mixtures of cord and adult blood cell-types as a means to further assess the reliability of UniLIFE. Using independent Illumina 450k data of 4 sorted cord blood and 6 sorted adult blood cell-subtypes, we generated such mixtures, revealing that UniLIFE can accurately estimate the known fractions (**Fig.4a**). We also generated in-silico WGBS mixtures of 5 cord blood and 10 adult blood cell-subtypes, which once again revealed excellent correlations and low RMSE values (**Fig.4b**), demonstrating that UniLIFE can accurately estimate fractions in complex mixtures, independent of underlying technology.

**Figure-4:**
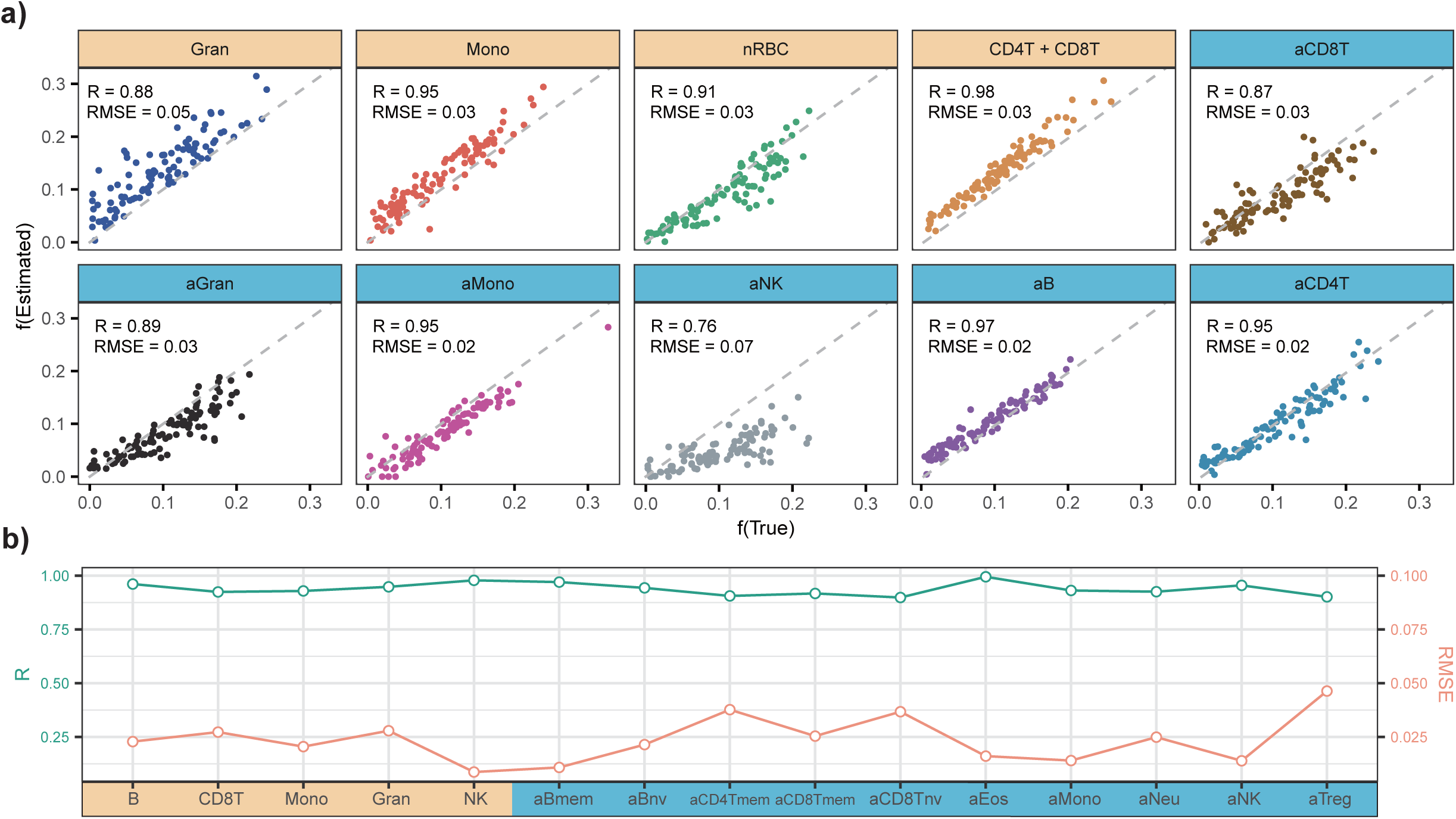
Validation of UniLIFE in blood of any age. **a)** For each of 4 cord blood cell types (orange) and 6 adult blood cell types (blue), scatterplots of true mixture weights (x-axis) vs estimated fractions (y-axis) for 100 in-silico mixtures of independent Illumina 450k DNAm profiles from Bell and DeGoede datasets. Each mixture is a linear combination of 10 immune cell sorted samples with weights drawn randomly from a uniform distribution. Pearson R-value and RMSE are given. **b)** As a), but now for 5 cord blood cell types (orange) and 10 adult blood cell types (blue) with WGBS DNAm profiles from IHEC. This dual-axis line chart illustrates the Pearson R-value (left Y-axis, green line) and RMSE (right Y-axis, orange line) for the true mixture weight and estimated fraction of each cell type across 100 in-silico mixtures. Each mixture is a linear combination of 15 WGBS immune cell sorted samples with weights drawn randomly from a uniform distribution.

### A map of immune-cell fraction variation throughout life reveals non-linear dynamics

To demonstrate practical utility of the 19 immune cell-type DNAm reference matrix, we assembled a large collection of Illumina DNAm datasets encompassing blood tissue from all main age-groups in a human life course, including preterm cord-blood (PreCB), cord-blood (CB), blood from newborns, 6 month old babies, children under the age of 10, adolescents (10-19 year olds), and all age groups up to a maximum of 94 (**Methods**). Some age-groups contained samples from various cohorts (**SI fig.S6a**), allowing batch effects to be assessed. We observed that batch effects were of a lower magnitude compared to the variation in cell-type fractions **(SI fig.S6b)**, allowing the dynamic change of these fractions during a human life course to be delineated. In support of this, we first note how cord-blood cell-type fractions were significantly higher in the PreCB, CB and neonate groups compared to adult samples where the fractions were close to zero (**Fig.5a**). The nRBC fraction was also distinctively higher in PreCB and in each of 4 independent CB datasets, as compared to adult cell-types, thus confirming that batch effects are smaller than the expected dynamic changes in cell-type fractions **(SI fig.S6c).** Conversely, adult immune cell-type fractions in PreCB, CB and neonates were very small or zero, subsequently displaying a progressive increase with age, with the exception of naïve lymphocyte fractions, which displayed a non-monotonic pattern, reaching maximum levels during the first 4 years of life (**Fig.5b**). The decreased naïve CD4T/CD8T fractions with increased age and the corresponding increase in memory fractions is well known ^46–48^. The rapid decrease in the cord-blood granulocyte fraction within the first 6 months of life is also well known ^31^, further demonstrating that our inferred patterns are mostly biological and not driven by batch effects. The non-monotonic pattern displayed by the naïve T-cell fractions was robust to the choice of DNAm datasets used, further attesting to the overall robustness of the dynamic patterns to batch effects (**SI fig.S6d**).

**Figure-5:**
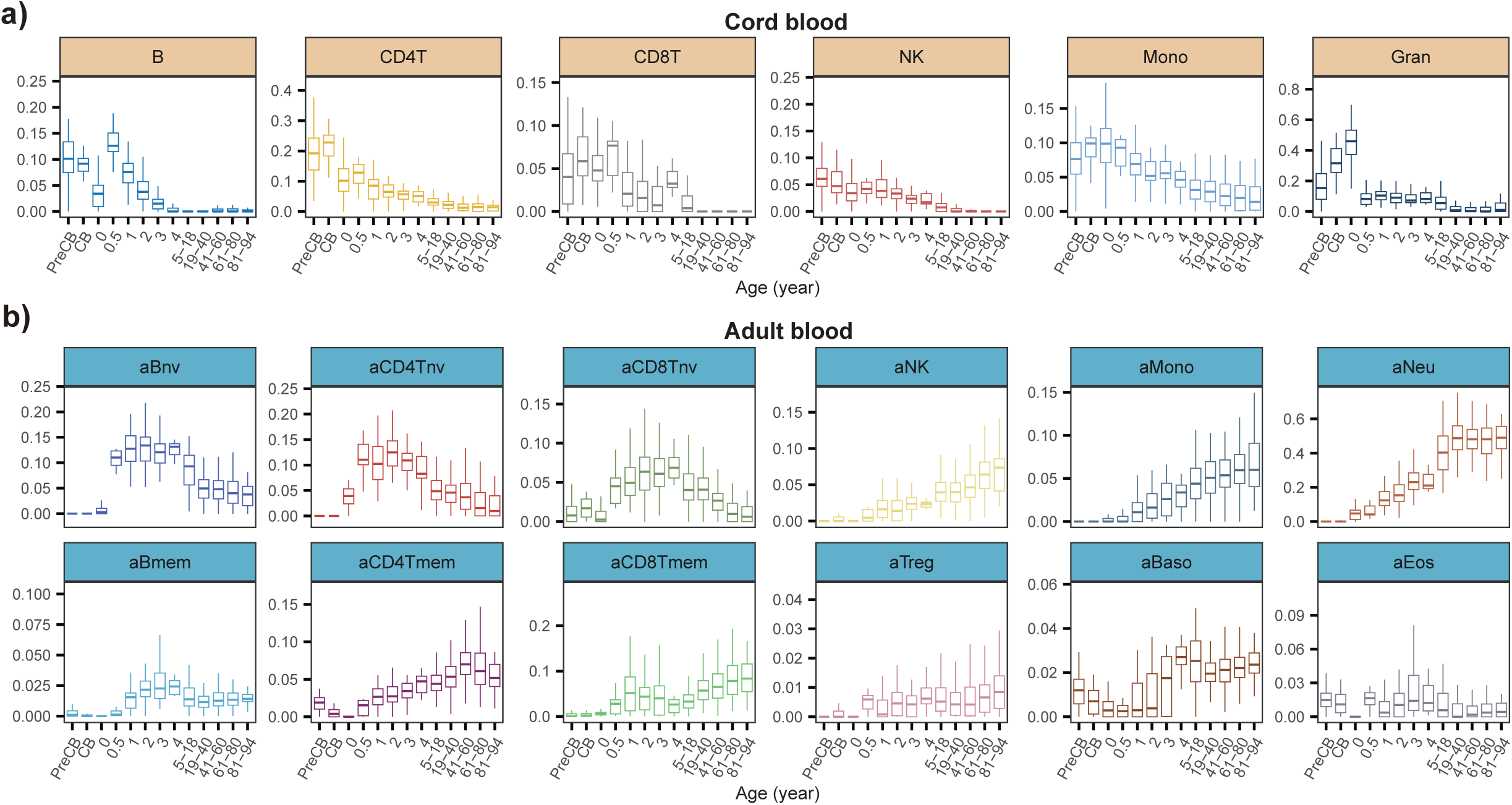
Immune-cell fraction variation throughout life reveals non-linear dynamics. **a)** Boxplots display the dynamic changes of 6 cord blood immune cell type fractions across all life stages. PreCB=cord blood from premature infants, and CB represents cord blood. After the age of 5, life stages are divided into age groups, including 5-18 years, and 20-year intervals thereafter. **b)** As a), but for the 12 adult immune cell types.

In order to further validate these dynamic patterns, we compared the specific dynamics of experimentally measured immune-cell type fractions within the first 6 years of life as reported by the Generation R study (GenR) ^31^, to our DNAm-based estimates (**Methods**). First of all, we observed relatively good agreement between our DNAm-based estimated cell-type fractions with those derived by GenR using flow cytometry (**Fig.6a**). Second, the predicted dynamics from the DNAm-based estimates was well correlated with those derived from flow-cytometry (**Fig.6b-c**). For instance, the drop in innate granulocyte and monocyte fractions within the first 6 months of life, as predicted with our DNAm reference matrix, was also observed using experimental cell counts (**Fig.6b**). In the case of naïve lymphocytes, for which our DNAm reference matrix predicted a peak at 6 months, we also observed a corresponding peak with the published experimental counts in the GenR-study (**Fig.6c**). Thus, overall, this data supports the view that UniLIFE can accurately chart dynamic changes in cell-type fractions within the first few years of life, an age-group for which we previously did not have a means to quantify such fractions from a DNAm reference matrix. This demonstrates the added-value of our DNAm reference matrix and shows that it is valid for any age-group.

**Figure-6:**
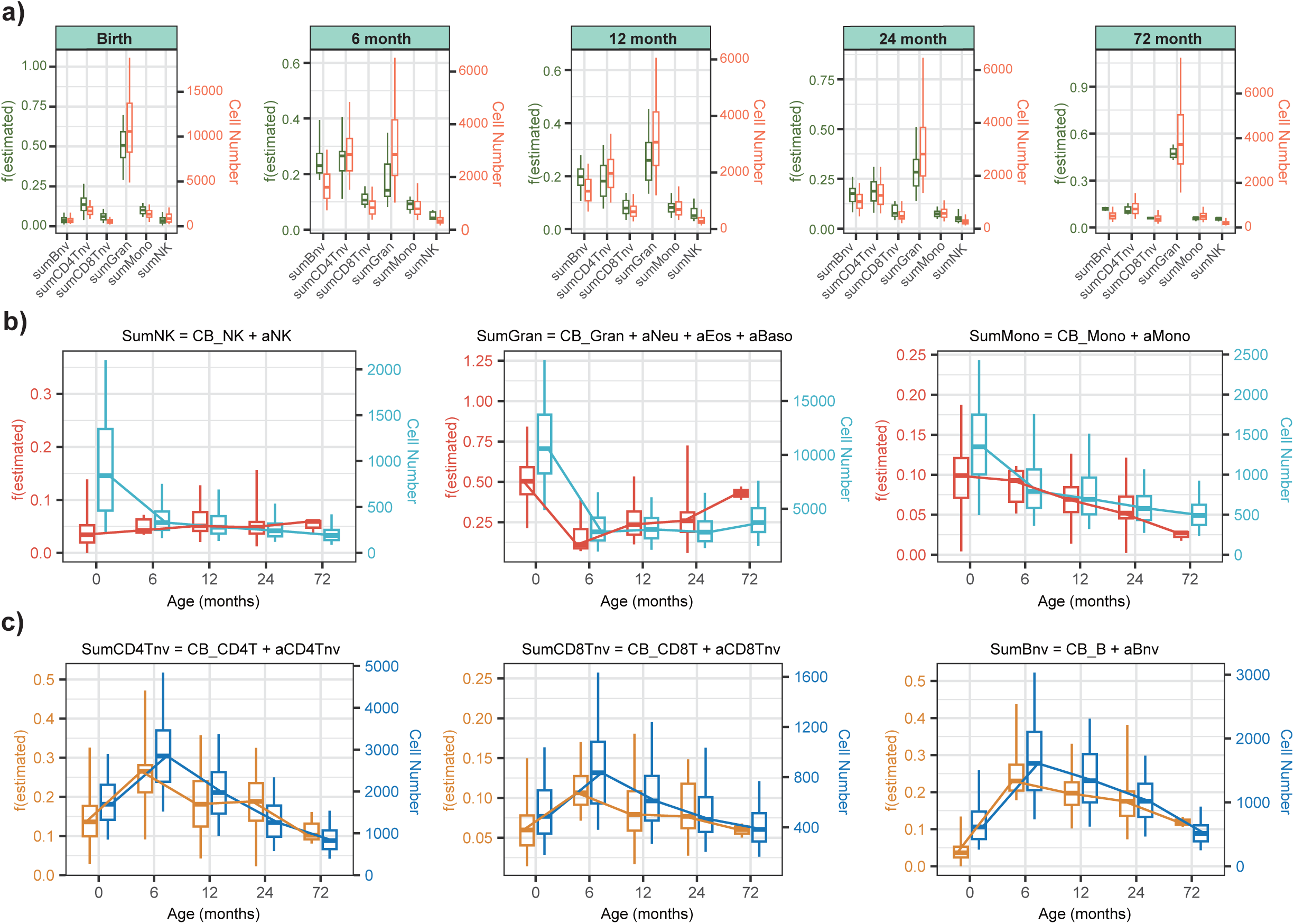
Comparison between DNAm-based estimates and cell counts reported by GenR. **a)** Boxplots compare the DNAm-based estimated fractions of 6 cell types (left y-axis) to experimental cell counts from GenR study (right y-axis) at 5 time points from birth to age 6. The f(estimated) (left y-axis) for each cell type is the sum of the fractions of cord blood type and the corresponding naive adult cell type (for lymphocytes) or the corresponding adult innate type (for myeloid cells), all fractions as estimated from the UniLIFE DNAm reference, i.e. sumBnv = CB_B-cell + aBnv, sumCD4Tnv = aCD4Tnv + CB_CD4T, sumCD8Tnv = aCD8Tnv + CB_CD8T, sumGran = CB_Gran + aNeu + aBaso + aEos, sumMono = CB_Mono + aMono, sumNK = CB_NK + NK. Cell Number (right y-axis) refers to the GenR experimental count of cell types. **b)** Boxplots compare the consistency of the variation patterns between the DNAm-based estimates and GenR experimental cell counts for three innate cell types measured in the GenR study from ages 0 to 6. The left y-axis, labeled f(estimated), corresponds to the DNAm-based fraction for sumNK, sumGran, and sumMono, while the right y-axis represents the corresponding experimental cell count. **c)** As b), but for 3 naïve immune cell types.

### Increased granulocyte to lymphocyte ratio in Down syndrome and preterm sepsis cases

Next, we asked if UniLIFE would be able to make concrete predictions about how immune cell-type proportions change in relation to diseases or conditions that affect infants. First, we considered Down syndrome (DS), since DS patients exhibit widespread immune system dysregulation ^49^. Specifically, using complete blood counts, it has been verified that neonates with DS generally exhibit neutrophilia and high nRBC counts including polycythemia ^50^. By applying UniLIFE to a HM450k blood DNAm dataset of neonates, encompassing 10 DS cases and 5 non-trisomy controls, we validated the increased cord blood granulocyte and nRBC fractions in DS neonates (**Fig.7a**). In contrast, cord blood CD4T, CD8T and B-cell fractions were lower in DS cases (**Fig.7a**), consistent with flow-cytometric analyses ^51^. To see if these changes persist in older DS cases, we estimated immune cell fractions in independent DNAm datasets of toddlers ^52^ and adults with and without DS ^53^. Granulocyte fractions in the older DS cases were no longer different from controls (**Fig.7a, SI fig.S7**), consistent with previous reports ^49^. However, the naïve CD4T, CD8T and B-cell fractions remained lower in adult DS cases, with the corresponding memory fractions being higher (**Fig.7a, SI fig.S7**). This is consistent with the higher risk of disease and lower life expectancy of DS patients.

**Figure-7:**
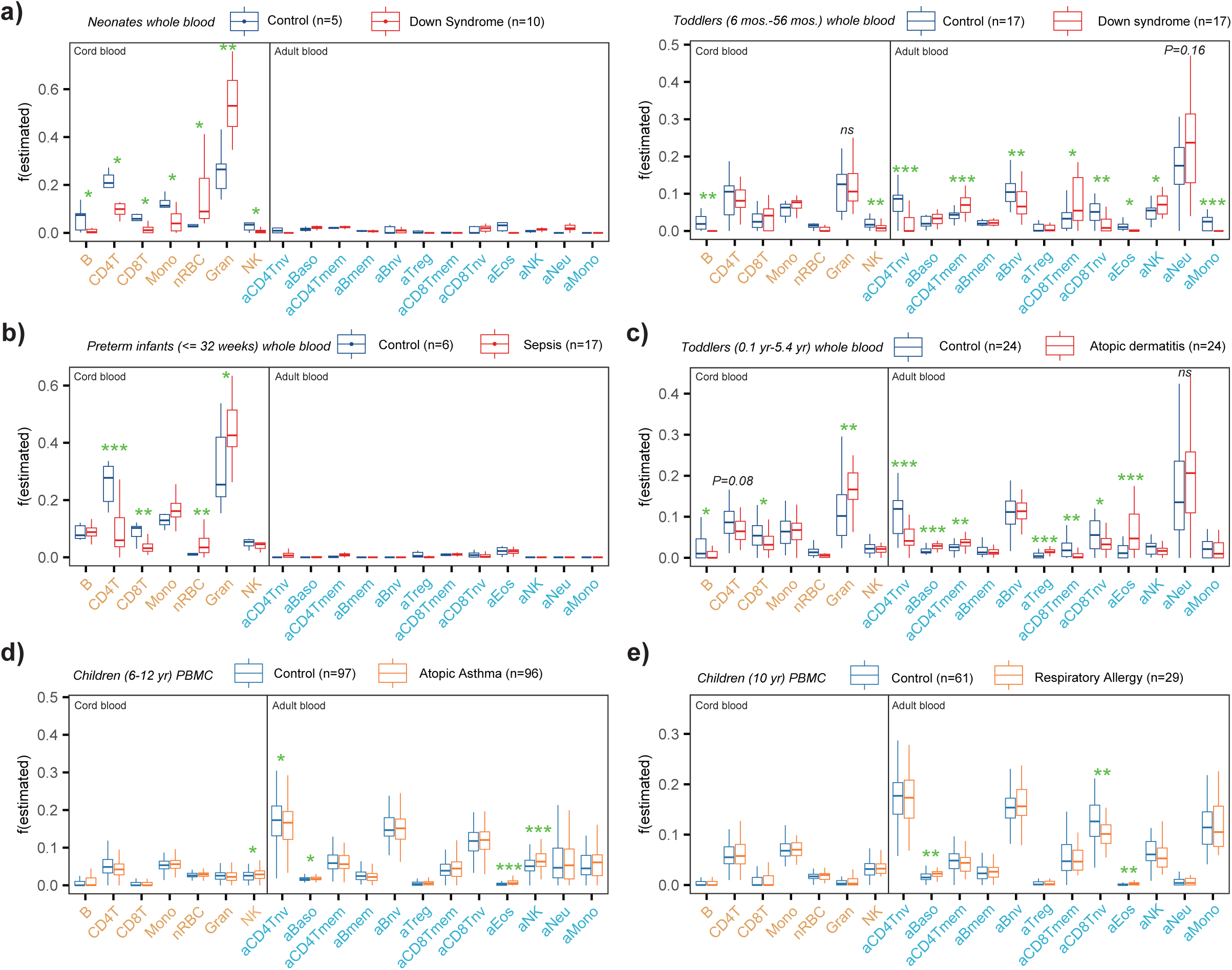
Validation of biological immune cell fraction variation. **a)** Boxplot displaying the estimated immune cell type fractions of neonates (left) and toddlers (right) with and without Down syndrome. Significance levels are derived from the Wilcoxon rank-sum test, with the symbols *** representing P < 0.001, ** representing P < 0.01, and * representing P < 0.05. **b)** As a) but for a DNAm dataset profiling whole blood of preterm infants with and without sepsis. **c)** As a) but for a DNAm dataset profiling whole blood from toddlers with and without atopic dermatitis. **d)** As a) but for a DNAm dataset profiling peripheral blood mononuclear cells of children with and without asthma. **e)** As a) but for a DNAm dataset profiling peripheral blood mononuclear cells of children with and without respiratory allergy.

Next, we applied UniLIFE to an EPIC cord blood DNAm dataset of 23 preterm infants, of which 17 developed sepsis ^54^. We observed how the granulocyte and nRBC fractions were significantly increased in sepsis cases compared to controls, whilst the T-cell fractions correspondingly decreased (**Fig.7b**), consistent with previous reports ^55,56^.

Finally, we applied UniLIFE to estimate immune cell fractions in 3 DNAm datasets of toddlers and children, encompassing atopic dermatitis ^57^, atopic asthma ^58^ and respiratory allergy ^59^ cases and age-matched controls (**Methods**). Although eosinophil and basophil fractions were small, we were able to detect, in each of the 3 conditions, a significant increase of these fractions in cases vs controls (**Fig.7c-e, SI fig.S8**), consistent with the role that these immune cell-types play in these diseases ^60–63^. Thus, overall, these data demonstrate the ability of UniLIFE to make correct predictions about biological immune-cell fraction variations in infants and children.

### UniLIFE infers DNAm changes in MHC-genes that predate the onset of type-1 diabetes

We next showcase how UniLIFE could be used in practice to infer differentially methylated cytosines (DMCs) in a longitudinal prospective study. We focused on an Illumina 450k/EPIC DNAm dataset, which profiled 395 blood samples from infants and children, at multiple timepoints during the natural progression of islet autoimmunity (IA) to type 1 diabetes (T1D) (the DAISY cohort) (**Fig.8a**) ^64^. This is an excellent case control study to test UniLIFE on, since prospective T1D cases and controls were matched for all confounders, including age, sex and technical factors (**Methods**). Using UniLIFE, we estimated the 19 immune cell-type fractions for each sample, which revealed that some fractions (e.g. the adult CD8+ T-cell memory fraction) displayed different dynamic profiles between cases and controls (**Fig.8b**). Specifically, whilst the adult CD8+ T-cell memory fraction increased with the timepoint of sampling in controls, it did not display a corresponding increase in the prospective T1D cases. It follows that cell-type heterogeneity, if not adjusted for, could negatively influence the selection of DMCs associated with T1D. To explore this, we ran linear models to find T1D-DMCs, adjusting for the 19 immune cell fractions in addition to other confounders that included sex and technical factors (chip and position) (**Methods**). We also adjusted for timepoint, in order to find T1D-DMCs that are consistently differentially methylated irrespective of whether the sample was taken before or after IA. Of note, this was done, because including an interaction term between disease status and timepoint to find T1D-DMCs where the effect size varies with the time to T1D diagnosis, did not reveal any significant associations, in line with previous findings from the DAISY study ^64^. Without an interaction term, we detected 3669 T1D-DMCs at an FDR<0.05 (**Fig.8c**). Using missMethyl ^65^, we found these to be enriched (FDR<0.0001) for over 40% of the genes in the KEGG type-1 diabetes mellitus pathway (**Fig.8d**). Adjusting for only 12 adult immune cell-types, or not adjusting for CTH, did not reveal any significant enrichment, despite the much larger number of T1D-DMCs (**SI fig.S9**), an indication that improper adjustment or no adjustment for CTH, substantially increases the false positive rate. By selecting the top ranked 3669 T1D-DMCs (thus matched for the number obtained with UniLIFE), we also observed enrichment of the KEGG T1D pathway, albeit weaker than when adjusting with UniLIFE (**Fig.8d**). Of note, the 19 enriched genes in the T1D KEGG pathway, included members of the MHC-class II (*HLA-DQB1, HLA-DQA1/2, HLA-DRB1/5, HLA-DMB, HLA-DOB*), the antigen presenting cell marker *CD86* and genes encoding autoantibodies (*GAD1* and *PTPRN2).* The list also included MHC-class I genes (*HLA-A/B/C/G/F*) and the cytotoxic CD8+ T-cell marker *FASLG* (**Fig.8e**) ^66^. Unlike the previous DAISY study, we were able to validate our T1D-DMCs (**Fig.8f**) in an independent DNAm dataset of sorted monocytes from 50 monozygotic adult twin pairs discordant for T1D ^67^. The validation in monocytes, but not in sorted B-cells or T-cells, suggests that the innate immune system may play an important role in the progression of IA to T1D, as recently discussed ^66^. Thus, overall, these results support the view that UniLIFE can aid the identification and interpretation of DMCs in longitudinal prospective EWASs performed in infants and children.

**Figure-8:**
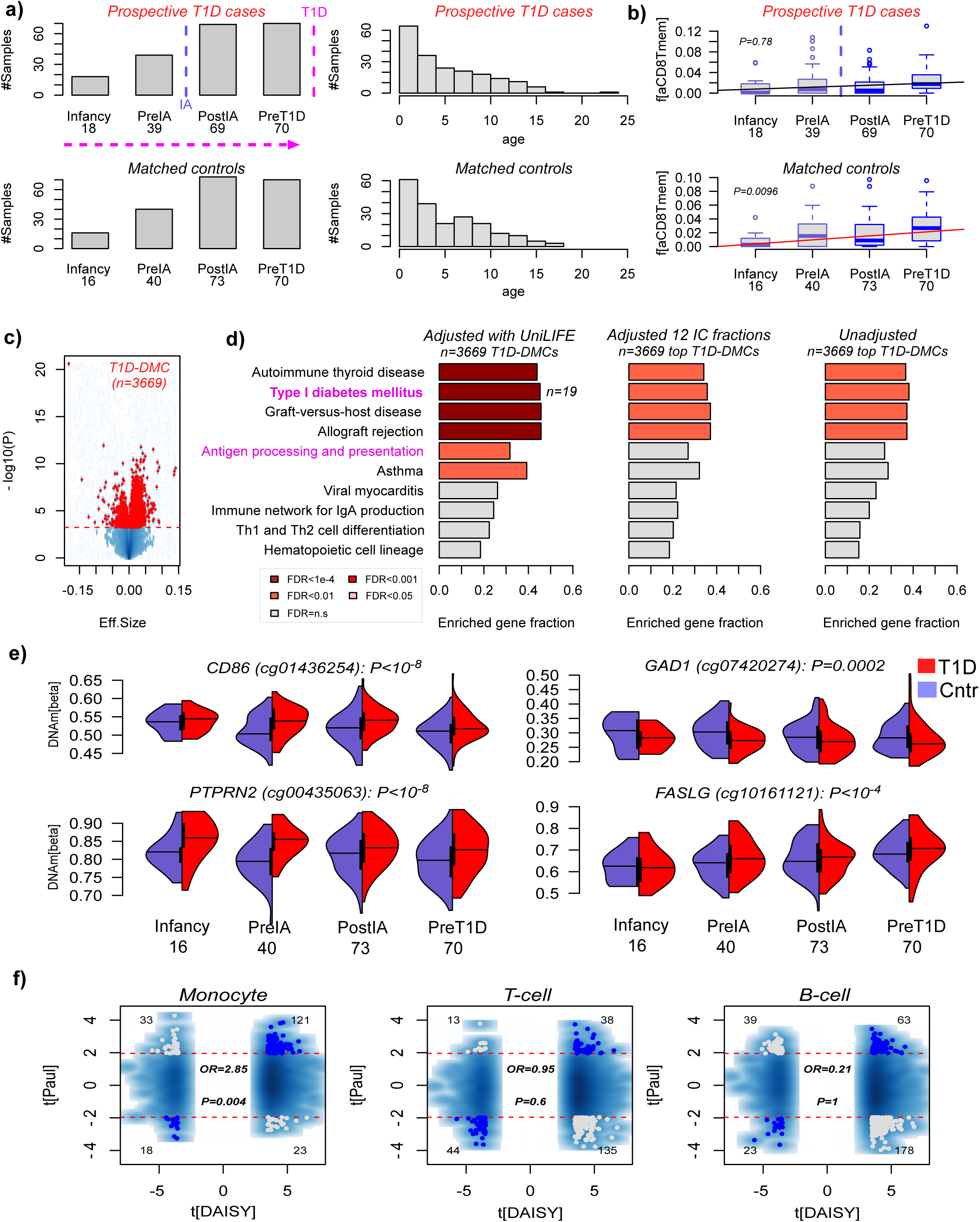
UniLIFE identifies differential DNAm in T1D pathway before T1D diagnosis. **a)** Left barplots compare the number of T1D (top) and control (bottom) samples across 4 distinct timepoints preceding T1D diagnosis in the cases. IA=islet autoimmunity. Right barplots display histograms of the samples according to age. **b)** Boxplot displays the estimated fraction of adult CD8+ memory T-cells as a function of timepoint in T1D cases (top) and controls (bottom). P-value is from a linear regression adjusted for age, sex, array-type, chip-position and chip. **c)** Volcano plot of T1D-DMCs as identified using a linear model that adjusted for array-type, chip, chip-position, sex, age, timepoint and 19 immune cell fractions as estimated using UniLIFE. **d)** Barplots depict the fraction of KEGG pathway genes that were found enriched among the T1D-DMCs derived with the model in c), using fractions estimated with UniLIFE (left), 12 adult immune cell fractions (middle) or no-adjustment (right). The numbers of T1D-DMCs in the latter two cases have been matched to the number obtained using UniLIFE. **e)** Violin plots of DNAm profiles of 4 T1D-DMCs mapping to KEGG T1D-pathway members, stratified by prospective disease status, as a function of timepoint. P-value is obtained from the same linear model. **f)** Validation of the UniLIFE inferred T1D-DMCs in a monozygotic adult twin study discordant for T1D status (Paul et al), where samples are sorted monocytes, B-cells or T-cells from approximately 50 twin pairs. The x-axis labels the t-statistics of association with T1D as in the discovery (DAISY) cohort, whilst the y-axis labels the corresponding t-statistics in the validation cohort (Paul et al). In each panel we count the number of T1D-DMCs that validate at a nominal P<0.05 threshold, or those that display inconsistency. The odds ratio (OR) and P-value from a one-tailed fisher’s exact test is given.

### Robustness of the UniLIFE reference panel to missing EPICv2 probes

Finally, given that the UniLIFE reference matrix was built from joint 450k-EPICv1 probes, it is important to assess if inferred cell-type fractions are robust to probes that may be missing on the recently released EPICv2 beadarray ^68^. From the 1906 CpGs in UniLIFE, 81 are missing or have been flagged in the EPICv2 release. Hence, to simulate performance in the EPICv2 setting, we re-estimated cell-type fractions removing these 81 probes. Across multiple different DNAm datasets, we observed very strong agreement between the newly estimated immune cell fractions and those estimated with the full repertoire of UniLIFE CpGs. This indicates that inference is very robust to removal of these 81 probes (**SI fig.S10**), and that UniLIFE would be applicable to new upcoming EPICv2 datasets.

## Discussion

Here we have advanced the field, by contributing a DNAm reference panel that works for blood tissue of any age. Although it was built using only sorted samples from cord and adult blood cell-types, these can be viewed as extremes in a continuum that includes blood from infants, children and adolescents. A key insight supporting this interpretation was derived by studying the principal components of variation of the merged DNAm matrix: this PCA revealed that adult naïve lymphocytes are more similar to their cord-blood counterparts than they are to adult (age-matched) mature lymphocytes, indicating that we have successful identified a maturity axis of DNAm variation. Importantly, this same axis of variation could also discriminate, albeit with a lower effect size, cord blood from adult innate immune cell-types, including granulocytes, monocytes and NK-cells. Thus, this same maturity axis of DNAm variation is simultaneously informative of age. As such, UniLIFE is able to chart the dynamic landscape of major immune cell-type proportions during the whole human life course.

We have extensively validated UniLIFE on (i) independent sorted cord and adult blood cell-types, (ii) on unsorted samples with matched experimental cell-counts and (iii) in-silico mixtures, encompassing two independent technologies (WGBS and Illumina EPIC/450k). Most importantly, UniLIFE makes predictions of dynamic immune cell type compositional changes within the first months and years of life that recapitulate those derived with experimental cell-counts in the GenR study. For instance, that naïve lymphocyte fractions reach a maximum at 6 months, remaining stable for the first 2 years of life, subsequently declining with age, as the corresponding memory fractions increase, is entirely consistent with those previously reported in a large study featuring repeatedly assessed cell counts ^31^. Further validating UniLIFE in infants and children, we have shown how we can correctly predict well-established dynamic cell-type compositional changes in a wide range of diseases and conditions, including Down syndrome, atopic dermatitis, asthma and respiratory allergy. For instance, we confirmed an increased granulocyte and nRBC fraction in neonates with DS compared to non-trisomy controls, but that this increase is no longer observed in children and adult DS cases. In contrast, naïve T and B-cell fractions, which are distinctively reduced in DS neonates, remain lower in children and adult DS cases compared to age-matched controls, which may underpin their lower life expectancy. Thus, the fact that UniLIFE can correctly predict these dynamic patterns for immune cell fractions during the first years of life, including adolescence, not only demonstrates the reliability of the estimated fractions, but also demonstrates its practical utility beyond what could have been achieved with a cord or adult-blood only DNAm reference matrix.

We have also showcased how UniLIFE can be applied to an EWAS performed in infants and children, to identify T1D-associated DMCs that predate the diagnosis of T1D. Specifically, our analysis revealed how specific immune cell-fractions (e.g. adult CD8+ memory cells) display different dynamics in prospective T1D cases compared to age-matched controls, indicating that proper adjustment for CTH could be critical. In line with this, our supervised analysis adjusting for 19 immune cell type fraction led to a stronger statistical enrichment for genes implicated in the T1D-pathway, despite a smaller number of T1D-DMCs, suggesting that improper adjustment for CTH using lower cellular resolution or not performing any adjustment leads to a larger number of false positives, consistent with previous reports on how CTH can inflate the false positive rate ^7,8^. The fact that using UniLIFE we could prospectively identify T1D-DMCs mapping to 19 genes in the T1D-pathway is highly significant, specially given that this was not reported by the original DAISY study ^64^. Moreover, unlike the original DAISY study, we were able to validate the pattern of differential methylation in association with T1D in an independent adult monozygotic twin study discordant for T1D status ^67^. Interestingly, the validation was seen in sorted monocytes, but not in B or T-cells, which could reflect the prospective nature of our T1D-DMCs, as it has been recently argued that innate immune cells play a more important role in the stages before T1D diagnosis ^66^. Although the monozygotic twin study predominantly comprises adults with diagnosed T1D (i.e. samples are postdiagnosis), it is plausible that DNAm changes in monocytes that predispose to T1D would still be present after diagnosis. Although this will require further investigation, our data clearly indicates how UniLIFE can lead to novel biological results and hypotheses.

It is important to end with a number of limitations. Although we have extensively validated our DNAm reference matrix, it would be desirable to have validated it in more independent sorted nRBC samples. We note that the sorted nRBC samples used in the construction of the DNAm reference matrix were more dispersed in the PCA plot, possibly reflecting subtle contamination with white blood cell-types ^43,69^. Indeed, nRBCs are known to interact with white blood cells, forming doublets that can result in cross-contamination when using conventional isolation techniques such as FACS or MACS ^43,69^. Thus, future improved iterations of DNAm reference matrices like UniLIFE would benefit from larger collections of sorted cord blood cell-types of higher purity, so as to increase the reliability and accuracy of the estimated fractions.

It is also worth pointing out that our UNILIFE DNAm reference matrix was built using joint 450k-EPICv1 probes. As such, further improvements could be envisaged if all sorted cord and adult blood data could have been generated with the same and more comprehensive EPIC platform. Such future attempts are very likely to use the recently released EPICv2 platform ^68^. With regard to this, we note that although approximately 17% of EPICv1 probes are not present on EPICv2, less than 5% (81 probes) of the 1906 CpGs in our UNILIFE reference panel are missing or have been flagged as unreliable from the EPICv2 array. As shown by us previously ^70^, and demonstrated again here, the inference of cell-type fractions from a DNAm reference matrix is very robust to missing probes and putative errors, tolerating up to 20-30% of missing data/errors without compromising reliability. This is because of the multivariate nature of the inference procedure, which is tantamount to a ‘voting or wisdom of crowds algorithm’, so that as long as the majority of the entries in the reference matrix are present and reasonably accurate, the algorithm will converge on the correct solution. Consequently, UniLIFE should be straightforwardly applicable to EPICv2, and indeed, as shown here, it is even applicable to orthogonal technologies such as WGBS.

A final limitation worth stressing is that UniLIFE was built from 19 sorted cell-types selected with specific surface markers, which may not be 100% specific. Moreover, there are substantially more than 19 immune cell types in the human immune system (by some estimates several hundreds) ^71^, with the number of functionally distinct immune cell-types also varying depending on the age of the subject. As such, the estimated cell-type fractions from UniLIFE cannot fully adjust for all underlying cell-type heterogeneity. Looking forward, it will be important to characterize the most relevant immune-cell subtypes for aging and disease, guided by single-cell studies, and to build correspondingly higher resolution DNAm reference panels.

## Conclusions

In summary, this study contributes a novel epigenetic software tool to enable unified estimation of 19 immune-cell type fractions across a human life-course. It is applicable to blood of any age, including cord-blood from preterm infants and newborns, as well as blood from infants, children, adolescents and adults. We envisage that this resource will enhance and aid interpretation of cross-cohort and longitudinal EWAS that aim to better model developmental/time-varying epigenetic signals in immune cell-types, by better dissecting the confounding effect of cell-type heterogeneity. The estimated immune cell fractions from UniLIFE may also serve as predictive or prognostic biomarkers. The UniLIFE DNAm reference matrix has been incorporated into our EpiDISH R-package, which is freely available from https://www.bioconductor.org/packages/devel/bioc/html/EpiDISH.html or https://github.com/sjczheng/EpiDISH.

## Methods

### Processing of the cord and adult blood DNAm datasets used in reference construction

#### Human sorted cord blood DNA methylation datasets used for building reference

##### Bakulski^44^

The HM450k dataset was downloaded from the Bioconductor package FlowSorted.CordBlood.450k, which provides a raw RGchannelSet object containing all sample information. In R (version 4.4.0), the preprocessRaw() and ratioConvert() functions from the minfi package (version 1.49.1) were used to convert the Red and Green channels of this object into a matrix of Beta values. The getAnnotation() function of the minfi package was used to identify and remove 11,458 probes located on sex chromosomes, and the dropLociWithSnps() function was employed to exclude 17,541 probes with potential SNPs in the CpG site and SNPs in the single base extension site. Additionally, 26,569 probes identified as having cross-reactivity were also excluded. A total of 1,787 probes were excluded for failing overall detection with P-values > 0.01 in any sample. The BMIQ algorithm was applied to scale bias of Type 2 probes. The final analysis included data on 428,157 probes across 104 samples, including 15 whole blood, 12 granulocytes, 15 monocytes, 15 B cells, 14 NK cells, 15 CD4+ T cells, 14 CD8+ T cells, and 4 nRBCs (nucleated Red Blood Cells).

##### Gervin^25^

The HM450k dataset was downloaded from the R package *FlowSorted.CordBlood-Norway.450k* in Bioconductor, which also provides a RGchannelSet object. Sample information such as cell type and gender are also contained within this object. The same preprocessing pipeline was applied as above, which involved the exclusion of 17,541 SNP probes, 11,458 sex chromosome probes, 26,569 cross-reactive probes, and 1,066 probes with a P-value > 0.01 in any sample. Ultimately, this resulted in data for 428,878 probes across 77 samples (with each of the following cell types has 11 samples: B cells, CD4+ T cells, CD8+ T cells, granulocytes, monocytes, NK cells, and white blood cells). Type-2 bias was corrected using BMIQ. Notably, this dataset does not possess nRBCs.

##### de Goede_1^43^

The HM450k dataset was downloaded from the GEO data repository (GEO accession GSE68456). After reading the IDAT files into a RGchannelSet object using the read.metharray.exp() function from the minfi package, the same preprocessing pipeline was applied as mentioned above, which involved the exclusion of 17,541 SNP probes, 11,458 sex chromosome probes, 26,569 cross-reactive probes, and 2,070 probes with a P-value > 0.01 in any sample. Finally, this resulted in data for 427,874 probes across 62 samples. Type-2 bias was corrected using BMIQ. The authors employed two sorting protocols: "standard" and "stringent." Whole (CD3+) T cells, monocytes, and nRBCs were collected from five individuals using the standard sorting method; B cells, CD4 T cells, CD8 T cells, granulocytes, monocytes, NK cells, and nRBCs were obtained from seven individuals following the stringent sorting protocol. In total, there were 7 B cells, 7 CD4+ T cells, 6 CD8+ T cells, 7 granulocytes, 12 monocytes (5 from standard and 7 from stringent), 6 NK cells, 5 nRBCs (from the standard protocol), and 7 nRBCs (from the stringent protocol), as well as 5 T cells. They have demonstrated that the standard sorting protocol can lead to cross-contamination among T cells, monocytes, and nRBCs. Therefore, subsequent analyses should exclude these 15 samples obtained from the standard protocol and instead use only those acquired through the stringent sorting protocol.

##### Lin^45^

The EPIC dataset was separated from the R package FlowSorted.CordBloodCombined.450k in Bioconductor, as the original source was inaccessible. Within the obtained RGchannelSet object, Gervin et al. retained only the HM450k probes. Then the aforementioned preprocessing pipeline was applied, which involved the exclusion of 16,353 SNP probes, 10,446 sex chromosome probes, 24,823 cross-reactive probes, and 918 probes with a P-value > 0.01 in any sample. The final analysis included data on 400,553 probes across 83 samples, which comprised 14 granulocytes, 14 monocytes, 13 B cells, 14 NK cells, 14 CD4+ T cells, and 14 CD8+ T cells. Type-2 bias was corrected using BMIQ. This dataset does not possess nRBCs too.

### Processing of the cord and adult blood DNAm datasets used for validation

#### Human cord blood DNA methylation datasets used for validation

##### de Goede_2^72^

The HM450k sorted cord blood dataset was downloaded from the GEO data repository (GEO accession GSE82084). 36 samples in total: 5 each of T cells, monocytes, granulocytes and nRBCs from term births; and 5 T cells, 4 monocytes, 4 nRBCs and 3 granulocytes from preterm births. Minfi package was used to preprocess the raw IDAT files. Subsequently, we excluded 17,541 SNP probes, 11,458 sex chromosome probes, 26,569 cross-reactive probes, and 598 probes with a P-value greater than 0.01 in any sample. This resulted in data for 426,642 probes. Type-2 bias was corrected using BMIQ.

##### Salas^27^

The EPIC dataset was downloaded from the GEO data repository (GEO accession GSE180970). There are 12 umbilical cord blood artificial mixtures with six cell types, including CD8T, CD4T, NK, B cell, monocyte, and granulocyte. After obtaining the raw IDAT files, 790,868 probes remain under the same preprocessing workflow. Type-2 bias was corrected using BMIQ.

##### Jones^73^

The HM450k cord blood dataset was downloaded from the GEO data repository (GEO accession GSE127824). The dataset includes 24 human whole cord blood samples with matched FACS counts of seven cell types, including CD8T, CD4T, NK, B cell, nRBC, monocyte, and granulocyte. Minfi package was used to preprocess the raw IDAT files and there are 429346 probes remained. Type-2 bias was corrected using BMIQ.

##### Hernandez^74^

The HM450k cord blood dataset was downloaded from the GEO data repository (GEO accession GSE149572). The dataset includes 3 adult human blood samples and 20 human cord blood samples. Beta value matrix with p-values was downloaded, and after removing the probes described earlier, 428,055 probes remain. Type-2 bias was corrected using BMIQ. 3 adult samples were excluded from the subsequent analysis.

##### Puvvula^75^

The HM450k cord blood dataset was downloaded from the GEO data repository (GEO accession GSE269983). The dataset includes 55 human whole cord blood mononuclear cell samples, 68 fetal side placenta tissues and 69 maternal side placenta tissues. Minfi package was used to preprocess the raw IDAT files and there are 422967 probes remained. Type-2 bias was corrected using BMIQ. Only 55 cord blood samples were used for subsequent analysis.

##### Holland^76^

The HM450k cord blood dataset was downloaded from the GEO data repository (GEO accession GSE97628). The dataset includes 380 human whole cord blood samples. Minfi package was used to preprocess the raw IDAT files and there are 391442 probes remained. Type-2 bias was corrected using BMIQ.

##### Bell_1^77^

The HM450k cord blood dataset was downloaded from the GEO data repository (GEO accession GSE188949). This dataset includes 107 cord blood samples from preterm neonates, of which 14 samples are from neonates with Bronchopulmonary Dysplasia (BPD). Minfi package was used to preprocess the raw IDAT files and there are 429730 probes remained. Type-2 bias was corrected using BMIQ. 14 samples with BPD were excluded from the subsequent analysis.

#### Peripheral blood datasets of children used for validation

##### Barwick^78^

The HM450k peripheral blood leukocytes dataset from healthy children was downloaded from the GEO data repository (GEO accession GSE36064). This dataset contains 78 samples from children aged 1 to 16 years. Minfi package was used to preprocess the raw IDAT files and there are 428593 probes remained. Type-2 bias was corrected using BMIQ.

##### Quinn^79^

The EPIC venous blood dataset was downloaded from the GEO data repository (GEO accession GSE224573). Sample size at each timepoint was as follows: n = 64 at birth, n = 8 at six months, n= 35 at one year, n = 32 at two years, and n = 16 at three years. Minfi package was used to preprocess the raw IDAT files and there remained 790585 probes after QC. Type-2 bias was corrected using BMIQ.

##### Wang^80^

The HM450k dataset was downloaded from the GEO data repository (GEO accession GSE104812). Samples included 29 boys and 19 girls, and their ages ranged from 6 to 14 years. Minfi package was used to preprocess the raw IDAT files and there remained 429426 probes after QC. Type-2 bias was corrected using BMIQ.

#### Human adult whole and sorted blood DNA methylation datasets used for validation

##### Bell_2 ^81^

The raw IDAT files for 789 sorted immune-cell samples of the HM450k platform were downloaded from the GEO data repository (GEO accession GSE224807). Any probe with a P-value greater than 0.01 was excluded as well as SNP probes and sex chromosome probes, resulting in a total remaining count of 456,116 probes. Type-2 bias was corrected using BMIQ. Among these 789 samples, there are six leukocyte subtypes: monocyte, granulocyte, B cell, CD4+ T cell, CD8+ T cell, and natural killer cell, as well as 36 whole blood samples (removed in subsequent analysis).

Other adult blood datasets used for validation have been previously normalized and analyzed in Luo et al., including: (1) *GSE112618*: 6 EPIC whole blood samples with matched FACS counts of 7 adult immune cell types, including CD4T cell, CD8T cell, B cell, NK, Monocyte, Neutrophil, and Eosinophil; (2) *GSE167998*: 12 EPIC artificial mixtures composed of 10 adult immune cell types: naïve B cell, memory B cell, naïve CD4T cell, memory CD4T cell, CD8T cell, NK, Mono, Neu, Eos, and Regulatory T cell (Treg); (3) *GSE56581:* 1202 monocyte and 214 CD4 + T-cell 450k samples. (4) *BLUEPRINT:* 139 monocyte, 139 naïve CD4 + T-cell, and 139 neutrophil 450k samples from the same 139 individuals. (5) *E-MTAB-2145*: 6 CD4 + T-cell, 6 CD8 + T-cell, 6 B-cell, 6 neutrophil, and 6 monocyte 450k samples. (6) *GSE65097:* 15 neutrophil 450k samples. (7) *GSE50222*: 8 CD4 + T-cell 450k samples. (8) *GSE63499:* 12 neutrophil 450k samples. (9) *GSE71955:* 31 CD4 + T-cell and 31 CD8 + T-cell 450k samples. (10) *GSE71244:* 6 CD4 + T-cell, 5 CD8 + T-cell, 4 B-cell, and 5 monocyte 450k samples. (11) *GSE59250:* 71 CD4 + T-cell, 56 B-cell, and 28 monocyte 450k samples. (12) *GSE59065:* 99 CD4 + T-cell and 100 CD8 + T-cell 450k samples. (13) *EGAS00001001598:* 49 CD4 + T-cell, 50 B-cell, and 52 monocyte 450k samples. (14)

##### GSE35069

6 CD4 + T-cell, 6 CD8 + T-cell, 6 B-cell, 6 neutrophil, 6 monocyte, and 6 eosinophil 450k samples. (15) *IHEC*: 4 memory CD4 + T-cell, 2 T-regulatory, 2 naïve CD8 + T-cell, 2 Cord blood CD8T, 2 memory CD8 + T-cell, 2 Cord blood B-cell,5 naïve B-cell, 5 memory B-cell, 2 Cord blood neutrophil,10 neutrophil, 20 monocyte, 2 Cord blood monocyte, 2 eosinophil, 2 Cord blood natural killer cell,and 2 natural killer cell WGBS samples. (16) HPT□450k (GSE210254): 418 HM450k PB samples; (17) Zannas (GSE72680): 422 HM450k PB samples; (18) Johansson (GSE87571): 732 HM450k PB samples; (19) HNM (GSE40279): 656 HM450k PB samples.

### DNA methylation datasets used for biological validation

#### Lorente-Pozo et al ^54^

This EPIC whole blood dataset was downloaded from the GEO data repository (GEO accession GSE155952). Blood samples were from septic (n=17) and healthy (n=6) preterm infants. Minfi package was used to preprocess the raw IDAT files. A total of 789721 probes remained. Type-2 bias was corrected using BMIQ.

#### Henneman et al ^82^

This HM450k whole blood dataset was downloaded from the GEO data repository (GEO accession GSE107211). Blood samples were obtained from 10 newborns with Down syndrome and 5 age-matched non-trisomic newborns. Minfi package was used to preprocess the raw IDAT files. A total of 429322 probes remained. Type-2 bias was corrected using BMIQ.

#### Naumova et al^52^

This HM450k whole blood dataset was downloaded from the GEO data repository (GEO accession GSE174555). Blood samples were obtained from 17 children with Down syndrome and 17 typically developing children (age ranges from 6 months to 56 months). Minfi package was used to preprocess the raw IDAT files. A total of 417613 probes remained. Type-2 bias was corrected using BMIQ.

#### Bacalini et al^53^

This HM450k whole blood dataset was downloaded from the GEO data repository (GEO accession GSE52588). Whole blood samples were obtained from 29 Down syndrome patients (age ranges from 10 years to 43 years) and 58 healthy controls (age ranges from 9 years to 83 years). Minfi package was used to preprocess the raw IDAT files. A total of 409225 probes remained. Type-2 bias was corrected using BMIQ. To ensure age matching, subsequent analyses only included 24 Down syndrome samples and 31 healthy control samples, all between the ages of 18 and 50.

#### Chen et al^83^

This HM450k whole blood dataset was downloaded from the GEO data repository (GEO accession GSE152084). Peripheral blood samples were taken from a total of 48 individuals (24 AD patients and 24 controls, age ranges from 0.1 year to 5.4 years). Minfi package was used to preprocess the raw IDAT files. A total of 415890 probes remained. Type-2 bias was corrected using BMIQ.

#### Yang et al^58^

This HM450k peripheral blood mononuclear cell (PBMC) dataset was downloaded from the GEO data repository (GEO accession GSE40576). PBMCs from 97 atopic asthmatic and 97 nonatopic nonasthmatic children (age ranges from 6 years to 12 years). Minfi package was used to preprocess the raw IDAT files. A total of 428167 probes remained. Type-2 bias was corrected using BMIQ.

#### Langie et al^59^

This HM450k PBMC dataset was downloaded from the GEO data repository (GEO accession GSE110128). It contains 29 Respiratory Allergy samples and 61 healthy control samples from children approximately 10 years old. Minfi package was used to preprocess the raw IDAT files. A total of 427883 probes remained. Type-2 bias was corrected using BMIQ.

### Processing of DNAm datasets of type-1 diabetes

#### DAISY study ^64^

The Illumina DNAm dataset was downloaded from the GEO data repository (GEO accession GSE142512). This dataset contains 395 samples from 174 individuals. The ages of the individuals range from 6 months to 22 years, with the majority being under 10 years old. A total of 211 samples were sequenced using the EPIC array, and 184 samples were sequenced using the 450k array. Half of the individuals were eventually diagnosed with type 1 diabetes, but all samples were taken before diagnosis. Samples from different array types were preprocessed separately. In R (version 4.4.0), the preprocessRaw() and ratioConvert() functions from the minfi package (version 1.49.1) were used to convert the Red and Green channels of IDAT files into a matrix of Beta values. Considering that the authors removed 7 arrays with inconsistent gender records during the quality checks, the estimateSex function from the R package wateRmelon (version = 1.0) was used to test the consistency between estimated and recorded gender. It was confirmed that the 7 arrays had already been removed from the raw IDAT data. The getAnnotation() function of the minfi package was used to identify and remove probes located on sex chromosomes (450k = 11458, EPIC = 19298), and the dropLociWithSnps() function was employed to exclude probes with potential SNPs in the CpG site and SNPs in the single base extension site (450k = 17541, EPIC = 30345). Additionally, probes identified as having cross-reactivity were also excluded (450k = 26569, EPIC = 39605). Any beta value with a detection P-value greater than 0.01 was considered as ‘missing’, and any probe with a coverage-fraction less than 0.9 was discarded. Missing values were then imputed using impute.knn (k=5) of R package impute (version=1.72.3). Type-2 bias was corrected using BMIQ algorithm. Finally, the EPIC and 450k normalized datasets were merged, defining a DNAm matrix over 400,485 common probes and 395 samples derived from 174 subjects. Of these 395 samples, 196 were from 87 subjects who would later develop T1D, with 199 samples being derived from 87 age-matched controls. Both controls and prospective T1D cases were sampled multiple times, with 22 subjects sampled 4 times, 49 subjects sampled 3 times, 57 subjects sampled twice and 46 subjects sampled only once. Samples were collected at 4 timepoints, two timepoints before the diagnosis of islet autoimmunity (IA), and two timepoints after diagnosis of IA but before diagnosis of T1D. Age range was 6 months to 22 years.

#### Paul et al ^67^

This is an Illumina 450k DNAm dataset of sorted monocytes, B-cells and T-cells from monozygotic twin pairs discordant for T1D, available from European Genome-phenome Archive (EGA) with the accession code EGAS00001001598 (https://www.ebi.ac.uk/ega/studies/EGAS00001001598). We used the normalized DNAm dataset as described in the original publication ^67^, consisting of 3 separate DNAm data matrices defined over 406,365 probes and 100 B-cell samples (50 twin pairs), 98 T-cell samples (49 twin pairs) and 104 monocyte samples (52 twin pairs).

### Construction of the universal 19 immune cell-type DNAm reference matrix

First, we focused on Illumina DNAm datasets profiling sorted cord blood cell-types that include CD4+ T-cells (CD4T), CD8+ T-cells (CD8T), NK-cells, B-cells, Monocytes (Mono), Granulocytes (Gran) and nucleated red blood cells (nRBCs). In total, we obtained four cord-blood Illumina 450k DNAm studies: Goede ^43^, Gervin ^25^, Bakulski ^44^ and Lin ^45^. All 4 studies contained on the order of 10 samples for each of the lymphoid and myeloid subsets, but only Bakulski and Goede contributed nRBCs (n=4 and n=7, respectively). For quality control, we used hierarchical clustering over the top significant singular vectors to cluster samples in each study. In three studies (Gervin, Lin and Goede), samples clustered perfectly by cell-type, with the exception of intermixing of CD4T and CD8T cells in Gervin. We verified using supervised analysis that we could identify many significant DMCs with large fold changes between the CD4T and CD8T cells from Gervin, suggesting that the observed intermixing from the unsupervised analysis is not necessarily a reflection of low purity. In the remaining study (Bakulski), nRBCs clustered separately from all other cell-types, but these sometimes did not segregate perfectly from each other, suggesting low purity of some of the leukocyte subsets. Hence, from Bakulski et al, we decided to only take forward the nRBC subset. We then merged the normalized DNAm data from the 4 studies, and performed a SVD and hierarchical clustering over the significant components. Whilst samples clustered by cell-type according to the first two singular vectors, the hierarchical clustering revealed some batch effects, for instance, the NK-cells of Goede clustered separately from the NK-cells of Lin and Gervin. Based on this hierarchical clustering we thus removed samples that did not cluster as expected. We subsequently reapplied SVD and hierarchical clustering, which resulted in a clustering where samples segregated perfectly by cell-type, with the exception of NK and B-cells. However, this splitting up of NK and B-cells had not been observed in our previous HC iteration, suggesting that this is a more minor effect. We argued that these residual batch effects are small and would not affect our downstream supervised analysis where we identify DMCs between cell-types. Supporting this, a scatterplot of the merged samples over the top two singular vectors resulted in a perfect segregation of samples by cell-type. The resulting sample size distribution for each cell-type is as follows: 28 B-cells (8 Gervin, 7 Goede, 13 Lin), 29 CD4T-cells (8 Gervin, 7 Lin, 14 Lin), 20 CD8T-cells (6 Goede, 14 Lin), 24 Granulocytes (8 Gervin, 2 Goede, 14 Lin), 32 Monocytes (11 Gervin, 7 Goede, 14 Lin), 22 NK-cells (8 Gervin, 14 Lin), 11 nRBCs (4 Bakulski, 7 Goede). Thus, for each cell-type, samples are drawn from at least 2 cohorts, and these samples cluster together, suggesting that their DNAm patterns are robust to residual batch effects. Overall, our merged sorted cord blood DNAm dataset was defined over 397,217 CpG probes and 166 samples encompassing 7 cord blood cell-types.

Next, we integrated this merged cord blood DNAm dataset with the EPIC DNAm dataset from Salas et al ^27^ encompassing 54 samples from 12 adult immune cell-types, a dataset which we have previously normalized and analysed in Luo et al ^29^. To distinguish these adult immune cell-types from those in cord-blood, we add the prefix “a” to indicate they are from adults: aBnv (adult naïve B-cells), aBmem (adult memory B-cells), aCD4Tnv (adult naïve CD4T-cells), aCD4Tmem (adult memory CD4T-cells), aCD8Tnv (adult naïve CD8T-cells), aCD8Tmem (adult memory CD8+ T-cells), aTreg (adult T-regulatory cells), aNK (adult NK-cells), aNeu (adult Neutrophils), aMono (adult Monocytes), aEos (adult Eosinophils), aBaso (adult Basophils). Thus, this resulted in an overall integrated DNAm dataset defined over 375,506 CpGs and 220 sorted samples, encompassing 19 immune cell-types (7 from cord blood, 12 from adult blood). We performed a SVD/PCA on this overall integrated DNAm dataset to ensure that the top PCs/SVs capture biology instead of batch effects.

To build the DNAm reference matrix for 19 immune cell-types, we need to identify cell-type specific DNAm markers. To identify these cell-type specific markers we used the following procedure:

1. For each cell-type, we performed differential DNAm analysis using an empirical Bayes method (limma) ^84,85^, comparing the cell-type to all the rest, using a stringent FDR<0.001 threshold.
2. Demanding for (i) hypomethylated cell-type specific markers that the attained maximum value over samples of that cell-type is less than the minimum value across all other cell-types. Likewise for (ii) hypermethylated markers we demand that the minimum value for the cell-type is greater than the maximum over all other cell-types.

We note that the second criterion is critically important to ensure specificity for a given cell-type, because criterion-1 alone does not guarantee specificity for each cell-type. For each cell-type, we finally ranked (i) hypomethylated CpGs by the gap in DNAm between the maximum value attained in the cell-type of interest and the minimum value across all other cell-types, and (ii) hypermethylated CpGs by the gap in DNAm between the minimum value attained in the cell-type of interest and the maximum value across all other cell-types. Thus, for each cell-type, we obtained a number of cell-type specific hypomethylated and hypermethylated markers. Aiming for approximately 100 markers per cell-type, we next tailored the marker selection for each cell-type as follows: for aBaso, aCD4Tmem, aBmem, aTreg, aCD8Tmem, aEos, aNK, aNeu, and aMono, we selected the top 100 hypomethylated marker CpGs, since the number of valid hypomethylated marker CpGs was greater than 100 for these adult cell-types. For the adult naïve fractions aCD4Tnv and aBnv we selected the top 50 hypomethylated and the top 50 hypermethylated markers. For aCD8Tnv, there were insufficient numbers of hypermethylated markers, so we selected the top 82 hypomethylated and the top 18 hypermethylated ones. For the cord-blood cell-types, the numbers of hypomethylated markers was generally always less than 100, requiring an approach that balanced hypo and hypermethylated markers. Specifically, for B-cells and nRBCs, we were able to choose the top 50 hypo and top 50 hypermethylated markers. For the remaining 5 cord blood cell-types (CD4T, CD8T, Mono, Gran and NK), the total number of hypo and hypermethylated markers was less than 100. Hence, for these cell-types we decided to supplement the marker CpGs with the top-100 hypomethylated CpGs ranked by fold-change, and not by the specificity gap score. Finally, we required marker CpGs to also display a range of DNAm-values across all cell-types of at least 0.2. In total, this resulted in 1906 marker CpGs, with approximately 100 markers per cell-type. The final DNAm reference matrix defined over these 1906 cell-type specific marker CpGs and 19 immune-cell types was built by taken the median value over the samples of a given cell-type.

### Estimation of cell□type fractions

In all cases, given the 19 cell-type DNAm reference matrix for either the Illumina EPIC or 450k dataset, we estimated corresponding cell-type fractions using the EpiDISH Bioconductor R-package. Specifically, we ran the *epidish* function with “RPC” as the method and *maxit* = 500.

### In-silico mixture validation analysis

In our study, we constructed three types of in-silico mixtures using the sorted cord blood dataset and the sorted adult blood dataset to simulate whole cord blood, whole adult blood, and infant whole blood respectively. The simulation of cord blood utilized all samples from de Goede_2, which include 10 T-cells, 9 monocytes, 9 nRBCs, and 8 granulocytes. For the simulation of adult whole blood, we utilized adult samples from the IHEC WGBS dataset. Therefore, the in-silico mixtures were generated from 4 memory CD4+ T-cell, 2 T-regulatory, 2 naïve CD8+ T-cell, 2 memory CD8+ T-cell, 5 naïve B-cell, 5 memory B-cell, 10 neutrophils, 20 monocytes, 2 eosinophils, and 2 natural killer samples. To simulate infant whole blood, we combined sorted cord blood samples and sorted adult blood samples, and concurrently constructed Illumina 450k mixtures and WGBS mixtures. The construction of Illumina 450k in-silico mixtures used all samples from de Goede_2 and Bell datasets, while the WGBS mixtures used all samples from the IHEC dataset (sorted cord blood samples included 2 CD8T, 2 NK, 2 B cells, 2 Neu, and 2 Monocytes). For each type of mixture, we generated 100 samples, randomly selecting one sample from each immune-cell type and using random weights drawn from a uniform distribution to generate the linear combination. Performance was assessed using Pearson R-values and Root Mean Square Error (RMSE).

### Comparison to Generation R blood cell counts

Percentiles of blood cell counts for Generation R study have been published in table E2 of the respective publication ^31^. Specifically, for each age group (0 months, 6m, 14m, 25m and 72m), leukocyte count values for all major immune cell types were given for the 5%, 25%, 50%, 75% and 95% centiles, allowing us to generate comparative boxplots.

### Identification of T1D-DMCs in DAISY and monozygotic twin studies

Because the DAISY DNAm dataset was generated with two separate array-types (450k & EPIC), and PC1 from a PCA analysis correlated most strongly with array-type, it was necessary to adjust for array-type when searching for T1D-DMCs. In addition, we observed non-negligible technical variation associated with array position and chip. Although cases and controls were correctly randomized and matched for age, timepoint, sex, array-type, position and chip, we adjusted for all of these factors when searching for T1D-DMCs. Specifically, we ran a linear model with DNAm beta-values as the response variable against prospective disease status, adjusting for array-type, array position, chip, sex, age and timepoint. Of note, this selects for DMCs where differential DNAm predates development of T1D in a manner that is also independent of timepoint, i.e. whether samples were collected before or after IA-diagnosis. T1D-DMCs were declared at a significance level threshold of FDR< 0.05. To identify CpGs where the differential DNAm between cases and controls increases or decreases with timepoint, we ran a model with an interaction term between disease status and timepoint, which however did not result in any significant hits. In the case of the monozygotic twin study, for each of the 3 cell-types (monocytes, B-cells and T-cells), we ran paired t-tests to estimate t-statistics and P-values for each CpG in association with T1D-status.

## Availability and requirements

**Project name:** UniLIFE within EpiDISH

**Project home page:** https://bioconductor.org/packages/release/bioc/html/EpiDISH.html

**Operating system(s):** Platform independent

**Programming language:** R(>4.0.0)

**License:** GNU GPL v3

**Any restrictions to use by non-academics:** none

## Declarations

### Ethics approval and consent to participate

We only analyzed publicly available data.

### Consent for publication

Not applicable.

### Availability of data and materials

The following DNAm datasets analyzed here are publicly available from the NCBI GEO website under the accession numbers GSE68456 (de Goede_1), GSE82084 (de Goede_2), GSE180970 (Salas), GSE127824 (Jones), GSE149572 (Hernandez), GSE269983 (Puvvula), GSE97628 (Holland), GSE188949 (Bell_1), GSE36064 (Barwick), GSE224573 (Quinn), GSE104812 (Wang), GSE224807 (Bell_2), GSE112618, GSE167998, GSE56581, GSE65097, GSE50222, GSE63499, GSE71955, GSE71244, GSE59250, GSE59065, GSE35069, GSE155952 (Lorente-pozo), GSE107211 (Henneman), GSE174555 (Naumova), GSE52588 (Bacalini), GSE152084 (Chen), GSE40576 (Yang), GSE110128 (Langie), GSE40279 (HNM), GSE210254 (HPT-450k), GSE72680 (Zannas), GSE87571 (Johansson), GSE142512 (DAISY). The sorted dataset of Zilbauer et al. can be accessed at ArrayExpress under accession number E-MTAB-2145. BLUEPRINT data can be accessed at EUROPEAN GENOME-PHENOME ARCHIVE (EGA) under accession number EGAD00010000850. And the sorted dataset of Paul et al. is also available from the EGA under accession number EGAS00001001598. International Human Epigenome Consortium data can be accessed at the IHEC data portal (https://epigenomesportal.ca/ihec/).

### Authors contributions

XG performed the statistical and bioinformatic analyses. AET, MS and AN assisted with bioinformatic analyses and literature curation. SCZ updates and maintains EpiDISH. BTH, AET and CAMC conceived the study and wrote the article.

