## Supplementary Figures for "Unified high-resolution immune cell fraction estimation in blood tissue from birth to old age"

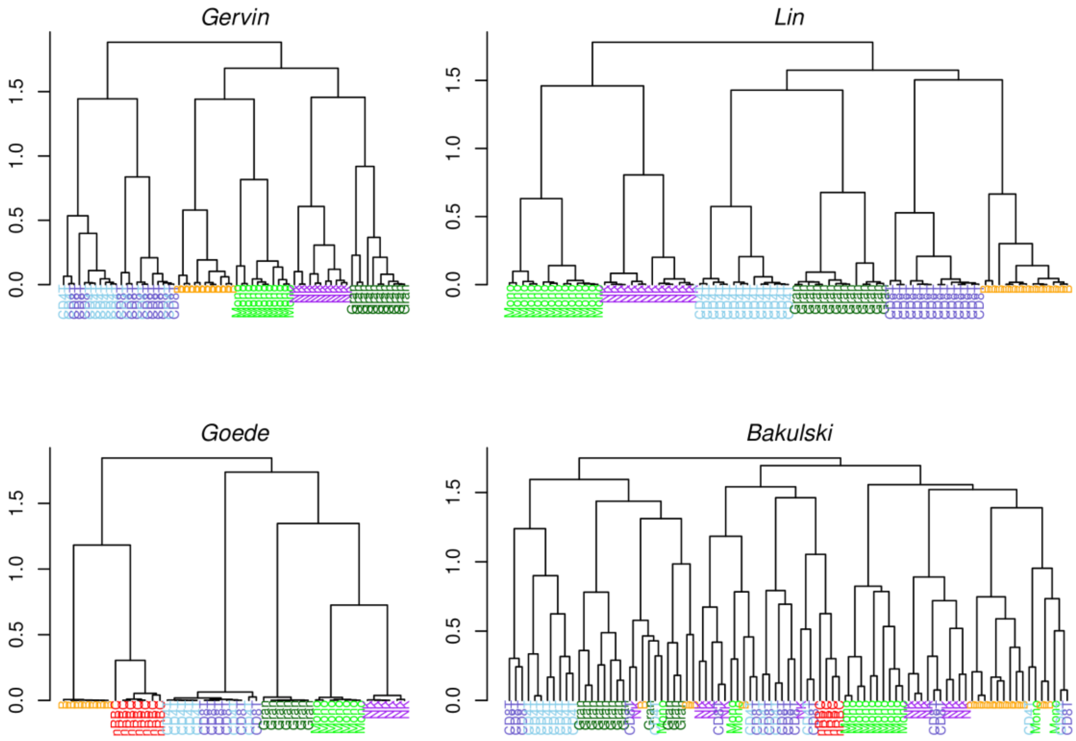


**SI fig.S1: Hierarchical clustering of the 4 cord-blood DNAm datasets prior to sample removal.** For each of the 4 studies we display the clustering dendrogram, with samples annotated by sorted cell-type. Color labels: skyblue=CD4T, darkblue=CD8T, orange=B, lightgreen=Monocytes, purple=NK-cells, darkgreen=Granulocytes.

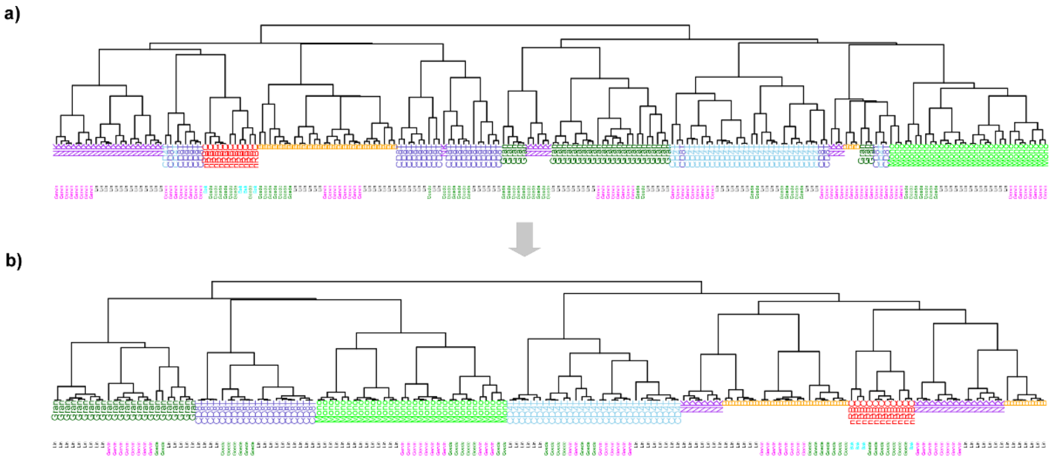

**SI fig.S2: Hierarchical clustering of the merged cord blood DNAm dataset. a)** Hierarchical clustering dendrogram of the merged samples from the 4 cord blood DNAm datasets, after removal of previous low purity samples. Sorted immune cell-types and study sorted sample is derived from is indicated below the dendrogram. Color labels for cell-types as in SI fig.S1. Color labels for studies: Magenta=Gervin, Black=Lin, Darkgreen=Goede, Cyan=Bakulski. **b)** As a) but after removing additional low purity samples, as inferred from a). The residual segregation of NK and B-cells is not observed in PCA-plots, and not observed in a), hence we conclude that this is not likely to cause confusion in downstream supervised analyses.

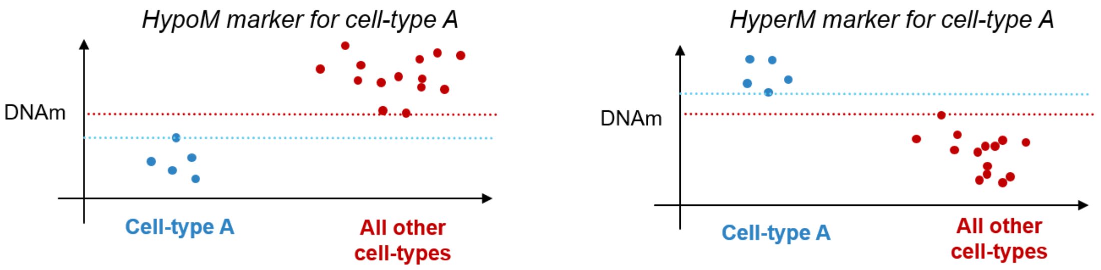


**SI fig.S3: Definition of Gap specificity score.** Diagrams illustrate hypothetical examples of cell-type specific hypomethylated (left) and hypermethylated (right) markers for a given cell-type "A" compared to all other cell-types, for which the gap specificity score is indicated as the difference between the blue and red dashed lines. For hypomethylated markers, we want the blue line to be lower than the red one, and vice-versa for hypermethylated markers. Features can be ranked according to this gap-specificity score, to ensure that markers are truly cell-type specific.

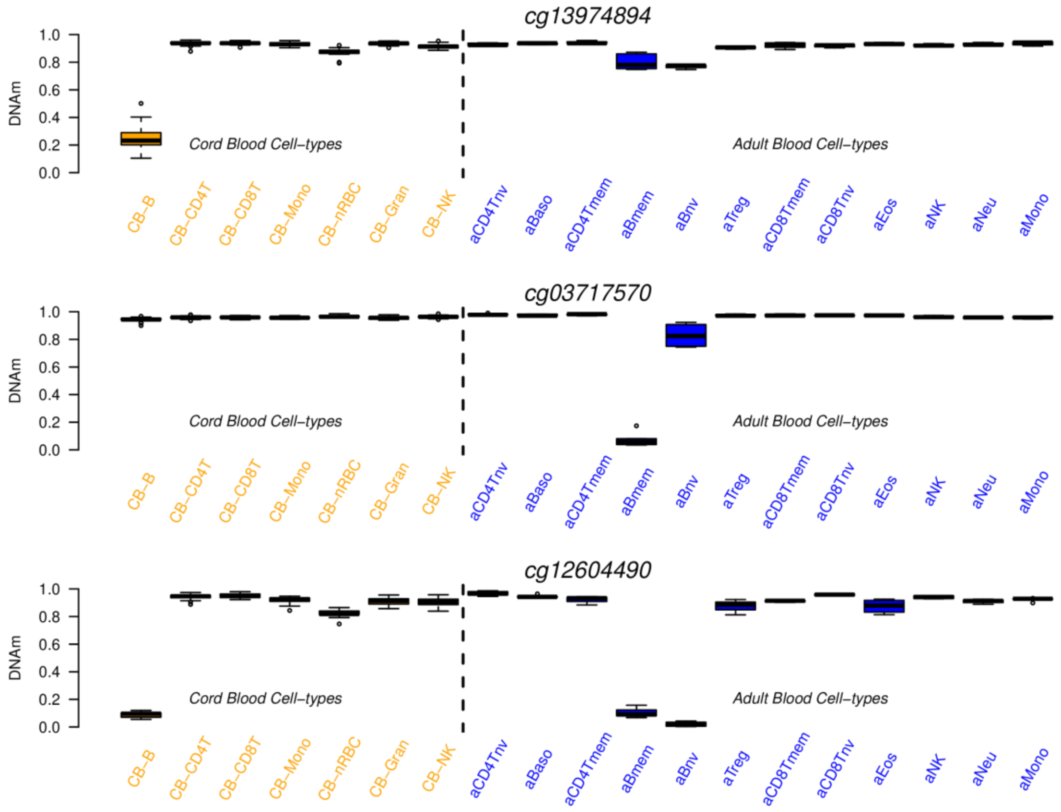


**SI fig.S4: Examples of selected markers for B-cell types.** Boxplots depicting the DNAm distributions of 3 selected markers across the 19 immune cell-types, the first six being from cord-blood (CB, orange labels) and the rest (12) being from adult blood (blue labels). The top marker is specific to cord-blood B-cells, the middle marker is specific for adult memory B-cells and the bottom marker is specific for all 3 B-cell types, although it is lowest for adult naïve B-cells. Whilst not strongly specific to naïve B-cells, to confidently distinguish these 3 B- cell types from each other it suffices to have the top 2 markers, with the bottom one distinguishing B-cells from the rest.

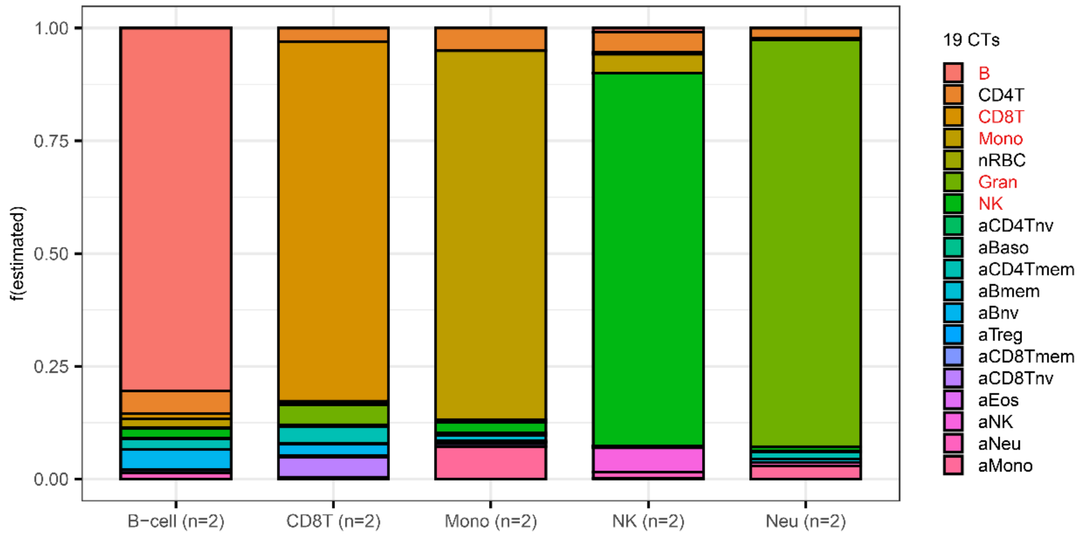


**SI fig.S5: Validation of UniLIFE DNAm reference in sorted WGBS cord blood samples.** The stacked bar chart displays the estimated cell type fractions of the UniLIFE DNAm reference matrix in five cord blood immune cell types. Each cell type has two samples, and the y-axis shows the average fraction over the two samples, attributed to each of the 19 immune cell subtypes indicated on the far right. It is evident that the largest section within each bar corresponds to the correct cell type.

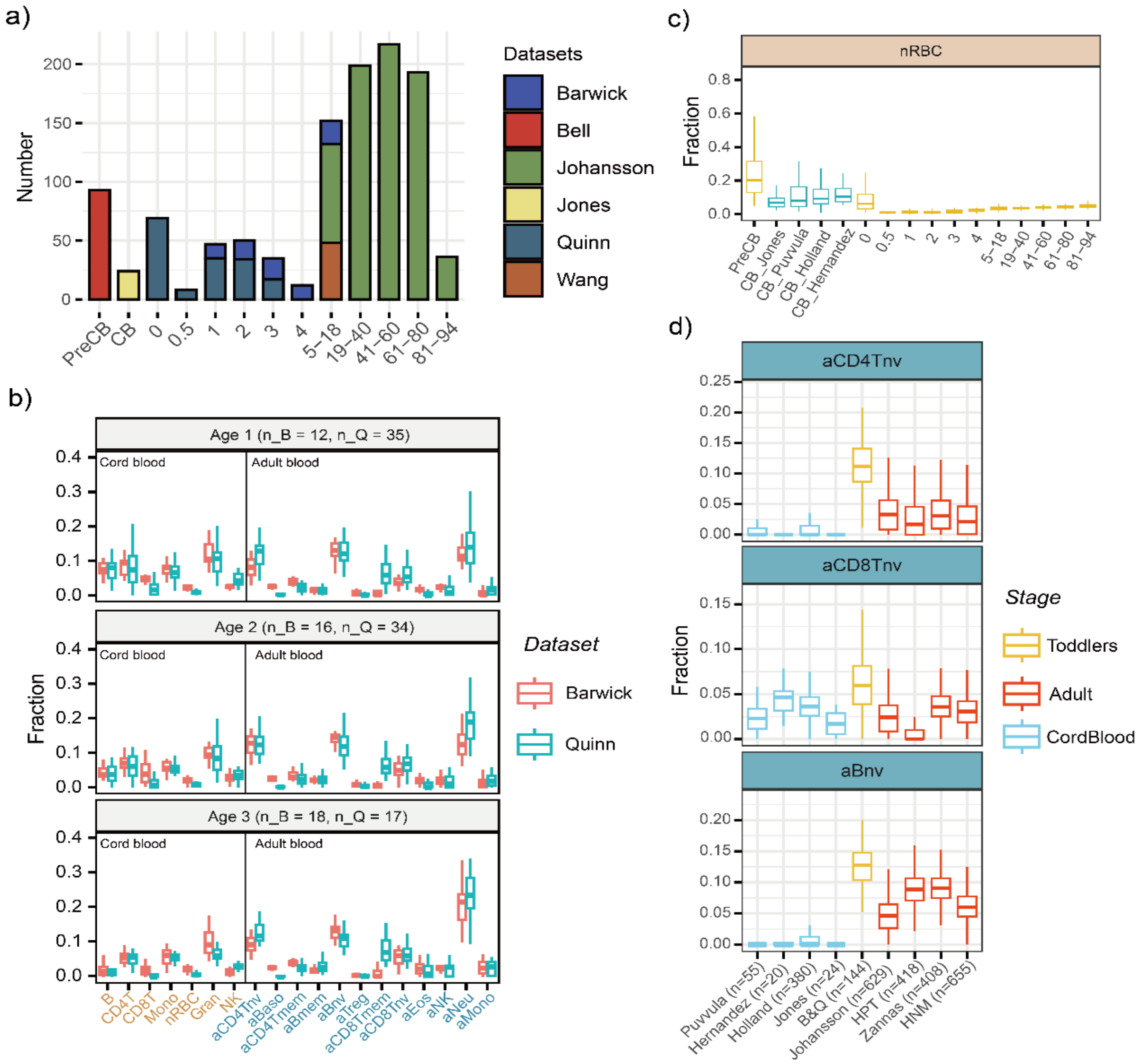


**SI fig.S6: Comparison of batch effects and age-related changes in cell type fractions.** **a)** The distribution of all datasets used in Main Figure 5 suggests potential batch effects between different datasets from various sources and age groups (PreCB: cord blood samples from premature infants, CB: corb blood samples from normal infants). **b)** A comparison of cell type fractions for each age from 1 to 3 years between the Barwick and Quinn datasets reveals that the fractions between datasets are similar. The number of samples included at each age for each dataset is provided (n_B: number in Barwick, n_Q: number in Quinn). Cord-blood cell-types labeled in orange. Adult-blood cell-types labeled in skyblue. **c)** Boxplot of nucleated red blood cell (nRBC) fractions across different age groups. In addition to the Jones CB dataset, three additional CB datasets (Puvvula, Holland, and Hernandez) were included to compare batch effects with true biological signals. **d)** Boxplots show that toddlers have the highest proportions of adult naive cell types, and this biological effect is significantly greater than the batch effect (three additional datasets of adults over 20 years old were included to demonstrate the robustness of the maximum for toddler group).

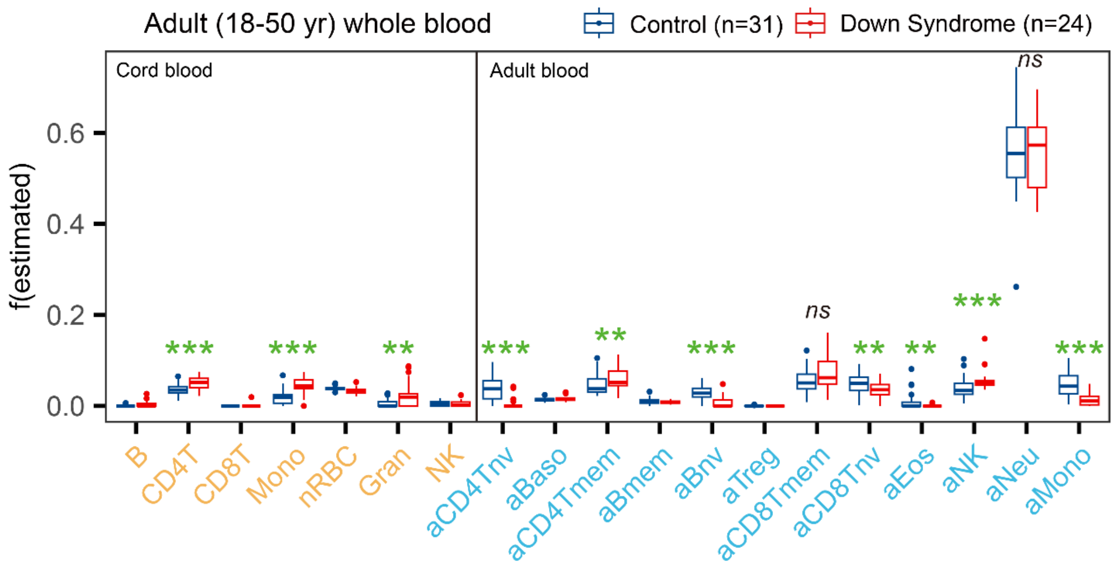


**SI fig.S7: Immune cell fraction variation in adult DS cases and controls.** Boxplots display the estimated 19 immune cell-type fractions in a DNAm dataset of 24 adult Down syndrome cases and 31 age-matched controls. Cord-blood cell-types labeled in orange. Adult-blood cell-types labeled in skyblue. Significance levels are derived from the Wilcoxon rank-sum test, with the symbols *** representing P < 0.001, ** representing P < 0.01, and * representing P < 0.05.

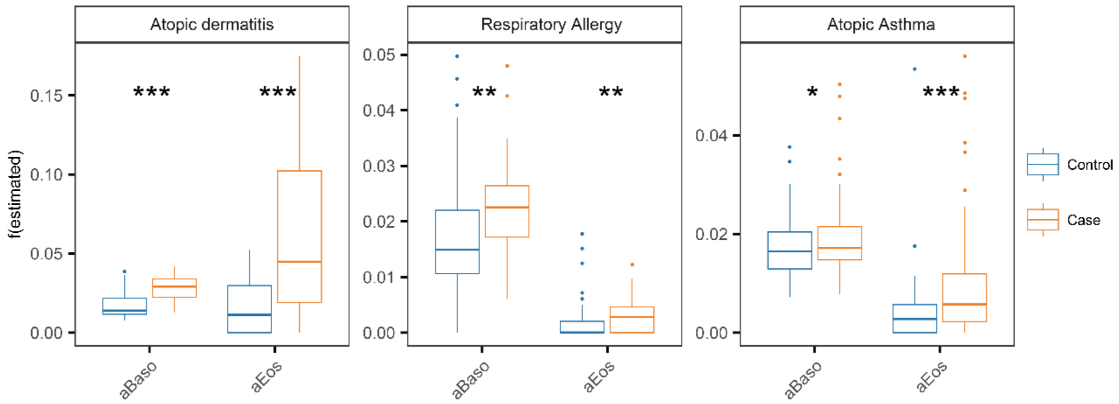


**SI fig.S8: Increased eosinophil and basophil fractions in dermatitis, allergy and asthma.** Boxplots display the estimated adult eosinophil and basophil fractions between cases and controls. From left to right, the cases are children with atopic dermatitis, respiratory allergy and atopic asthma, with controls being age-matched. Significance levels are derived from the Wilcoxon rank-sum test, with the symbols *** representing $P < 0.001$, ** representing $P < 0.01$, and * representing $P < 0.05$.

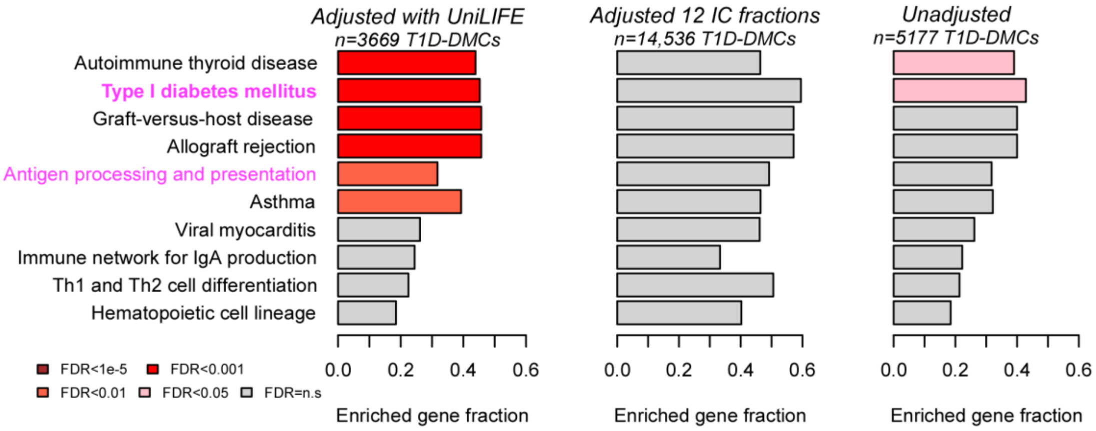


**SI fig.S9: Comparative enrichment analysis using common FDR threshold.** Left: The enriched fraction of genes (x-axis) of the top-10 enriched KEGG pathways among the 3669 T1D-DMCs (FDR<0.05) derived with UniLIFE in the DAISY prospective T1D study. Color of bars indicate statistical significance of enrichment (FDR) level, as shown. Middle & Right panels: As left, but now for the 14,536 T1D-DMCs (FDR<0.05) and 5,177 T1D-DMCs (FDR<0.05) derived using adjustment with 12 adult immune cell fractions or with no adjustment for cell-type heterogeneity.

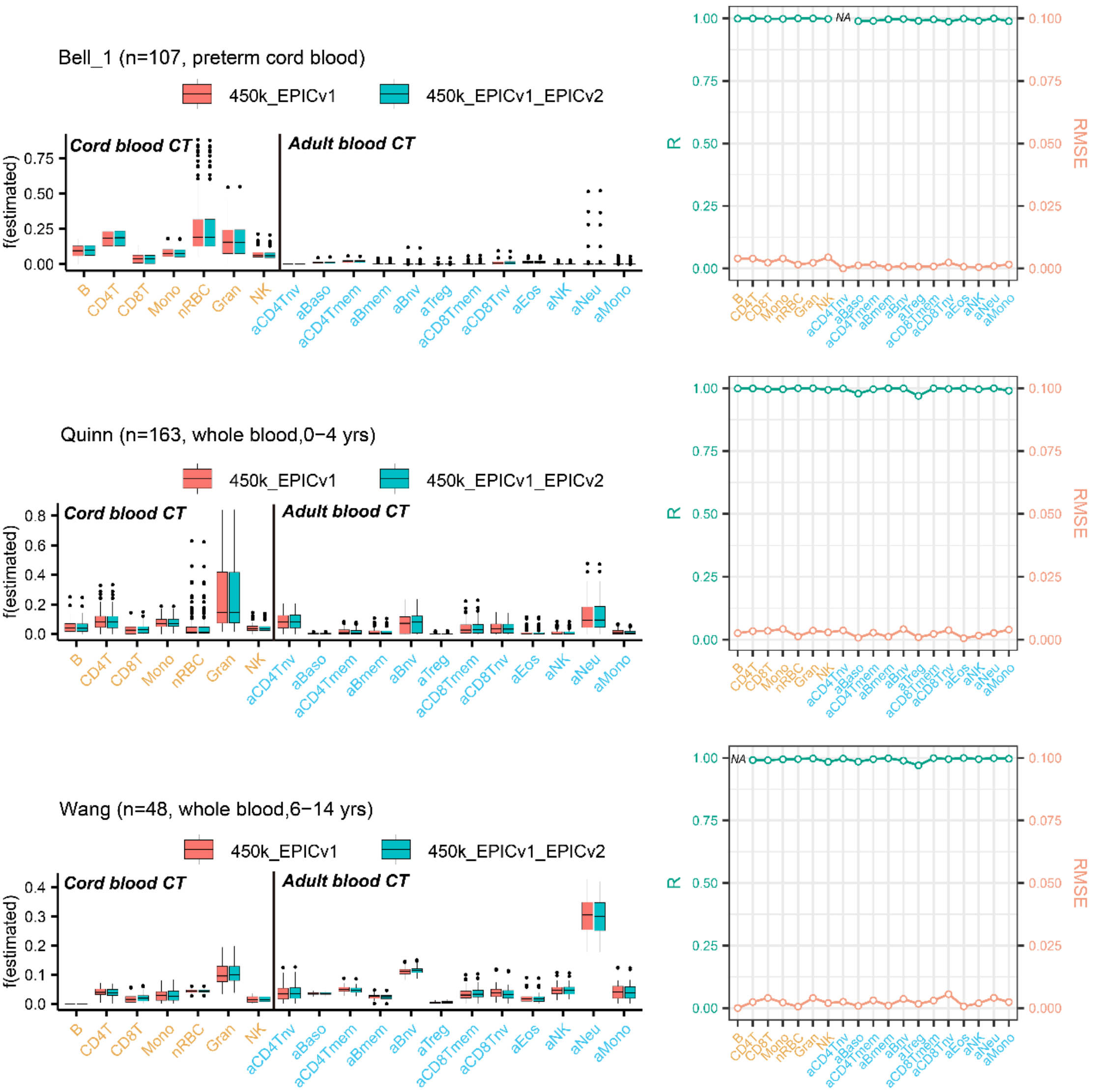


**SI fig.S10: Robustness of cell-type fraction estimation to missing probes on EPICv2.** Left panels: For each of 3 Illumina DNAm datasets we contrast boxplots of estimated cell-type fractions, as obtained with UniLIFE using all joint EPICv1-450k CpGs in the reference (red) and by removing those not present on EPICv2, i.e. using only joint EPICv1-EPICv2-450k probes (blue). Right panels: For each dataset, R-values and RMSEs for all cell-types. Cord-blood cell-types are labelled in orange, adult blood cell-types are labelled in skyblue. NA means that no R-value (Pearson Correlation Coefficient) is computable because all fractions were zero.
